# Laser Scanning Confocal printing, a new simple method for micro-device fabrication (LSCprint method)

**DOI:** 10.1101/2024.02.08.579477

**Authors:** D Megías, J Gómez, J Ortega, G Mata, M. Pérez-Martínez

**Author notes:** Equally contribution (D Megías, J Gómez).

## Abstract

Micro-devices and lab-on-a-chip technologies have revolutionized cell biology research, enabling a plethora of applications from single-cell sorting to organ-on-a-chip assays. However, their construction remains laborious, requiring specialized equipment and trained personnel, thereby restricting their accessibility to specialized laboratories. The conventional protocol for micro-device printing involves intricate steps, including master cast/mold production and device fabrication, leading to high costs and time consumption.

Here, we present a novel, simplified method utilizing a laser scanning confocal microscope (LSCM) and commercially available photosensitive resins. By using the UV or violet excitation laser lines of an LSCM, we eliminate the need for external suppliers and complex equipment, enabling any conventional cell biology laboratory to fabricate micro-devices swiftly and inexpensively.

Our method not only enhances the capabilities of standard confocal microscopes but also democratises microfluidic device fabrication, making it accessible to non-specialized laboratories. With minimal reagent consumption and high scalability, our approach offers a cost-effective solution for rapid prototyping and production of micro-devices, circumventing previous barriers to widespread adoption. Moreover, our method allows direct printing of micro-devices onto substrates, eliminating the need for molds and intermediate steps, thus facilitating greater design flexibility and accessibility for non-specialized laboratories.

## Introduction

In cell biology research, we can find many kinds of micro-devices and lab-on-a-chip successfully used for a wide variety of applications, including single-cell sorting^1-4^, cell cycle studies^2,5-7^, cell migration assays^8^, 3D growing cells scaffolds^9-13^, experiments for chemical reactions^14,15^ and even organ-on-a-chip assays^16,17^. However, their construction and development are still laborious, and they need highly trained staff and dedicated equipment. In conclusion, this kind of technology is usually restricted only to specialized laboratories.

The most widely spread protocol for micro-device printing can be sorted in two steps. The first one begins with a suitable design that leads to the production of a master mold^6,14,15,17-19^. The created mold have hole spaces that are filled with liquid resins or silicones which, after some chemical and development processes, will turn into solid, yielding a 3D structure with the pattern of interest^6,14,15,17-19^ (Figure 1a). This is a critical step that requires very particular equipment^18^ and frequently needs to be outsourced to specialized laboratories or commercial suppliers. In the latter case, this step has an associated fee, which varies depending on size and complexity of the design, but it also introduces an important delaying time in the fabrication of the final device^18^.

**Figure 1.**
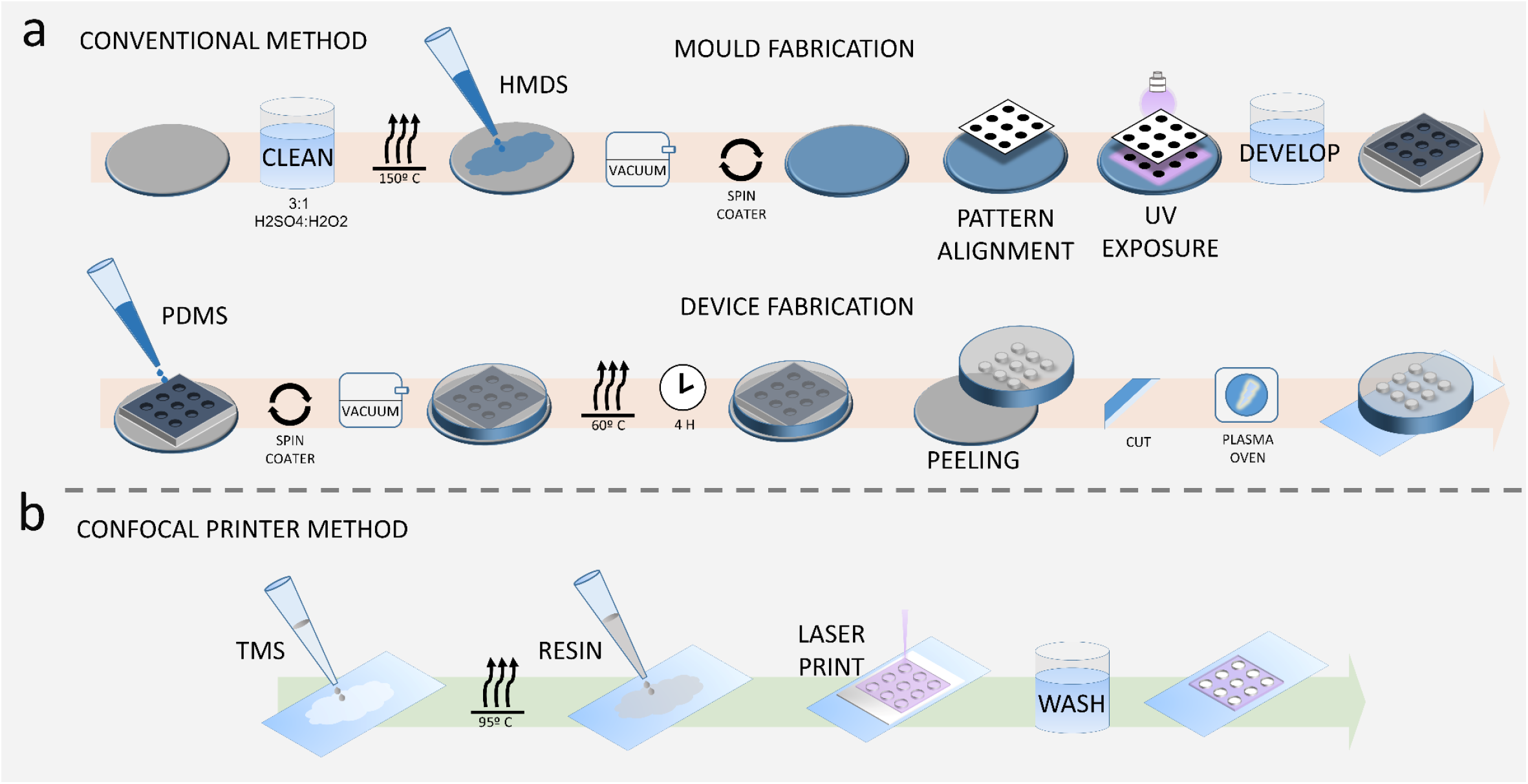
Comparison of micro-devices fabrication methods **a** Conventional fabrication method showing the process from mold fabrication to final device development. The steps include dedicated equipment for pattern transference and device fabrication, such as master mold fabrication machine, spin coater, heaters, vacuum chambers and harmful reagents. **b** Method based on confocal printing using a photosensitive resin. TMSPMA is used to increase device attachment, and washing steps remove any rest of un-polymerized resin.

Once the master mold is obtained, the second step of the micro-device fabrication protocol can start. From this point on, the protocol is complex and requires a laborious and long procedure. This includes the access to special reagents and the use of dedicated equipment such as a plasma oven, to bind the micro-device to the substrate (glass or plastic) where the final assay will take place^14,18,19^; a spin coater to generate a flat and uniform layer of resin or silicone^6,14,18,19^; a vacuum chamber used to degas the liquid resin (preventing the appearance of air bubbles into the final device)^6,14,18^, and optionally, an oven that triggers the polymerization reaction^6,14,15,18-19^ (Figure 1a). This is a critical step that requires very particular equipment^18^ (Figure 1a).

After the silicone polymerization, the jellified layer must be peeled off from the waffle. Then the portions required for the assay are manually cut and bound to the final substrate (usually a glass slide or a coverslip)^18^ (Figure 1a).

The protocol described above can be repeated as many times as needed to create a high number of micro-devices, but any variation or improvement of the original design will require to generate/order a new waffle mold and to re-start the whole process, hindering possible improvements and the best refinement of the mentioned devices. This could be a problem for non-specialised laboratories interested in this technology, for example biomedical groups, since a protocol so long and expensive results in an entrance barrier, difficult to surpass

The new alternative method that we propose here reduces the fabrication costs and time by eliminating steps and the necessity to contract any external supplier to produce master molds. This benefit is achieved by using accessible reagents and equipment. With the proposed simplified workflow, any conventional cell biology laboratory, without needing an extensive experience in the use of this technology, will be able to generate a wide variety and amount of its own micro-devices within the time frame of a few minutes to a few hours, depending on the complexity of the device.

This new micro-fabrication method avoids the need of the expensive and time-consuming mold production by using a laser scanning confocal microscope (LSCM), usually easily accessible to most of laboratories, and commercially available photosensitive resins, which are conventionally used in regular 3D printers and a few extra reagents.

In order to convert an LSCM into a micro-device printer, the instrument just needs to have two technical characteristics, quite widespread. It must have ultraviolet (360 nm) or violet (405nm) excitation laser lines, which are commonly used for regular fluorescence assays, and the capability of directing the laser to desired regions of illumination (ROIs) to enable the control of sensitization and hardening of the photo-resins. After the mentioned controlled light exposure, the final device is printed as a solid surface and, after some simple washing steps, it will be biocompatible and suitable for live-cell imaging experiments (Figure 1b)

This new method not only expands the possibilities of conventional confocal microscopes but also gives a real opportunity to non-specialized laboratories to create their own microfluidic devices.

Our proposed method is extremely affordable in terms of time and reagents, since a bottle of 200 ml of resin is not expensive (see materials section). Furthermore, we use volumes that are in the range of few microliters, so just one resin bottle may allow (depending on the surface to be covered) the production of more than 4,000 micro-devices, and may last for several years, depending on their use.

Some previous attempts have been reported to devise alternative fabrication methods^12,20-22^, including the use of circuits built on paper^23-27^, or using other methods, such as high-power laser stereo lithography or direct laser writing^10,13,28,29^. Again, the poor resolution^22^ or the need of special requirements and equipment^18,22,28^ have represented an important barrier, thus limiting the spread of the micro-fabrication technology.

The novel method described here can be used to create a mold for conventional production, helping this way to generate prototypes until the final design is achieved. But more importantly, it opens the possibility to directly print the final micro-device on the glass surface that will be used, skipping the step for the generation of the mold, without any intermediate steps. This new alternative gives more freedom for micro-device design and fabrication, making it widely accessible to non-specialized laboratories.

## RESULTS

### Generation of the micro-device

The method proposed for the elaboration of a photo-printed micro-device is illustrated in Figure 1b and can be summarized along the steps detailed below:

1. Increase the adherence of polymerized resin to the glass surface (when the surface is porous, such as in plastic, this part can be skipped) This step is performed by covering the glass surface with 3-(Trimethoxysilyl)propyl methacrylate (TMSPMA)^5,6^ for just a few seconds. After this step, the coverslip is placed on a heater plate at 95ºC for 30 seconds, so the excess of TMSPMA evaporates, and only the fraction attached to the glass remains. To further clean the rests of TMSPMA, and as an optional step, the coverslip may be rinsed with isopropanol 100%, with no negative effect over the adhesive capabilities of the TMSPMA layer.
2. Resin layer formation Pure resin can be placed directly onto the glass surface, but it would form a rounded drop, not a spread uniform layer. In order to get a thin and uniform layer, resin can be mixed in diethyl-ether in the proportion of one part of resin per three parts of diethyl-ether (1:3). Depending on the resin formulation, this proportion may vary. After this mix preparation, it must be spread all over the surface with the help of a pipette tip. In just a few seconds, most of the diethyl-ether evaporates, leaving a thin homogeneous layer of resin. The mix volume to add onto the surface depends on the area itself and the desired thickness; for example, for covering a 25 mm rounded coverslip, only 50-100 microliters of the mix are needed.
3. Confocal photo-printing process In the confocal microscope, the first step would be to focus the sample on the resin plane. This is usually achieved by capturing laser reflection with a non-violet laser, to find the interface between resin and surface. Next, you must draw/load the regions of illumination (ROIs) and run the acquisition process as in a common image capture, using a violet (405 nm) or ultraviolet (365 nm) laser. Depending on the laser power of the system, it could be useful to add additional acquisition steps, such as frame/line averaging, to ensure proper polymerization (Suppl. Material 01).
4. Washing step Once the photo-polymerization process is finished, the device must be washed in order to remove the non-polymerized resin. For this purpose, the micro-device must undergo three washing steps, one minute each, with xylol 100%, ethanol 100%, and distilled miliQ water, following that order. After that, the device can be air-dried with no negative effect over its resistance or capabilities.
5. Resin hardening Depending on the hardness required, an extra step of illumination of the now-clean full micro-device can be done by exposing it to a fluorescence UV lamp. This can be done using the microscope’s fluorescence lamp, or even with a violet laser pointer (405 nm).
6. Steps for bio-compatibility and sterilizing Place the glass with the device on a heater, 95ºC for 30 seconds, with the glass surface in contact with the heater, not the device surface. This helps to evaporate any remaining solvent. Let the coverslip be washed 3 times, 5 minutes each, in abundant Phosphate-buffered saline (PBS). Let the coverslip with the device dry. You can speed up this process using compressed air. Sterilize it with UV light for at least 30 minutes in a culture hood.
7. Cell seeding can be done as in a regular culture plate.

This protocol results in a micro-printed device firmly attached to a coverslip, and depending on your experiment, ready to be used.

For the design transference (3^rd^ step in the protocol), an accurately controlled illumination of the regions of interest (ROIs) must be achieved. Fortunately, nowadays most of the laser scanning confocal microscopes have user-friendly drawing tools to define these ROIs, so you can manually draw them or copy and paste customized illumination areas. These ROIs can be saved to reproduce the same micro-pattern in future experiments, or even to perform *in situ* modifications to the original design (Figure 1b).

### Micro-device sealing

For some experiments, the created device must be sealed^30-33^, since unlike conventional microfabrication, our method produces channels and reservoirs that are open (Figure 2a). The sealing can be easily done by filling the channels with melted paraffin, and keeping the resin surface as clean as possible (Figure 2b and 2c). Once it cools, paraffin will block the channels with a solid coating (Figure 2c), allowing to add new resin on top of the device and to polymerize it with a 405-laser pointer, or with the fluorescence lamp of the microscope using the proper UV filter, to generate a layer that will cover the whole device (Figure 2d).

**Figure 2.**
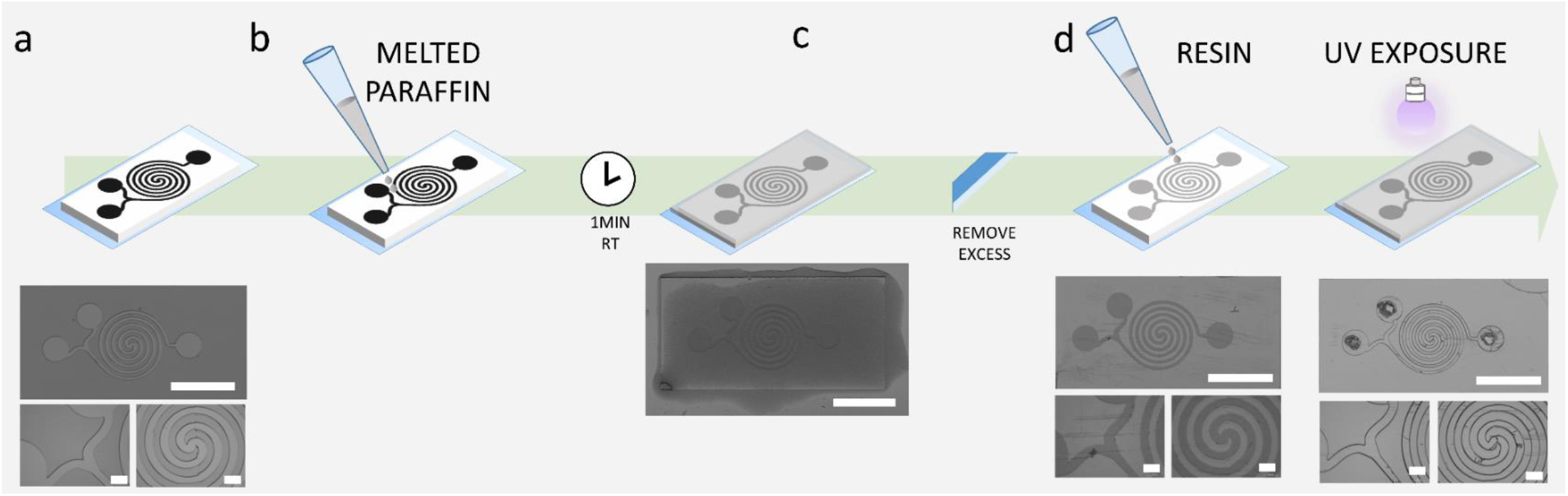
Sealing of a micro-device. **a** Printed pattern with open channels. **b** By adding melted paraffin open channels are reversibly closed. **c** After paraffin cooling, the excess of paraffin is removed with a razor. **d** Addition of fresh resin to coat the device and hardening with UV. Scale bars are 5 mm for complete device and 500 μm for detailed zooms. All images attached to schematic representations are brightfield images captured with a magnification glass.

If you want to avoid resin polymerization in some areas (for example, to insert inlets or outlets), you can just draw some black spots with a black marker under the glass surface before the polymerization step, and illuminate from beneath with the microscope’s fluorescence lamp or the 405-laser pointer, so the UV light will not pass through these black areas.

After this process, the paraffin that is filling the device’s channels can be removed by melting it with heat (60º C) and displacing it with air pressure (using a syringe, for example). After removing the paraffin, further cleaning of the device should be performed with distilled water.

Another possible approach is to print the pattern and then attach it to commercially available sticky plates, that will add inlets/outlets and/or main walls. The lids that these sticky plates provide work fine for open design patterns such as single cell traps, etc., but they are not suitable for device sealing such as in high pressure closed flow circuits (Figure 2).

### Photo printed micro-devices performance

Printing resolution depends on the confocal microscope used (laser power, illumination configuration, etc.), on the proportions of the resin and solvent used for the mixture, and on the thickness of the resin layer (the thicker the layer, the lower the resolution) (Figure 3). Based on this, different shapes and sizes are possible. By using the microscope described in Materials and Methods section, a maximum printing resolution of 3 microns can be achieved (Figure 3) on a resin with a thickness of 6.5 microns. In this sense, this new method is comparable to conventional methods in terms of resolution. This resolution should be more than enough for the majority of cellular assays^1,2,4-8,11-16,18-20,22,26,30,33^. Additionally, size comparison experiments between designed regions and printed ones showed little differences, with an average difference of 1.5 um, and a maximum difference of 3.4 um (this maximum corresponding to a designed region of 32 um). Further results are shown in Figure 3c.

**Figure 3.**
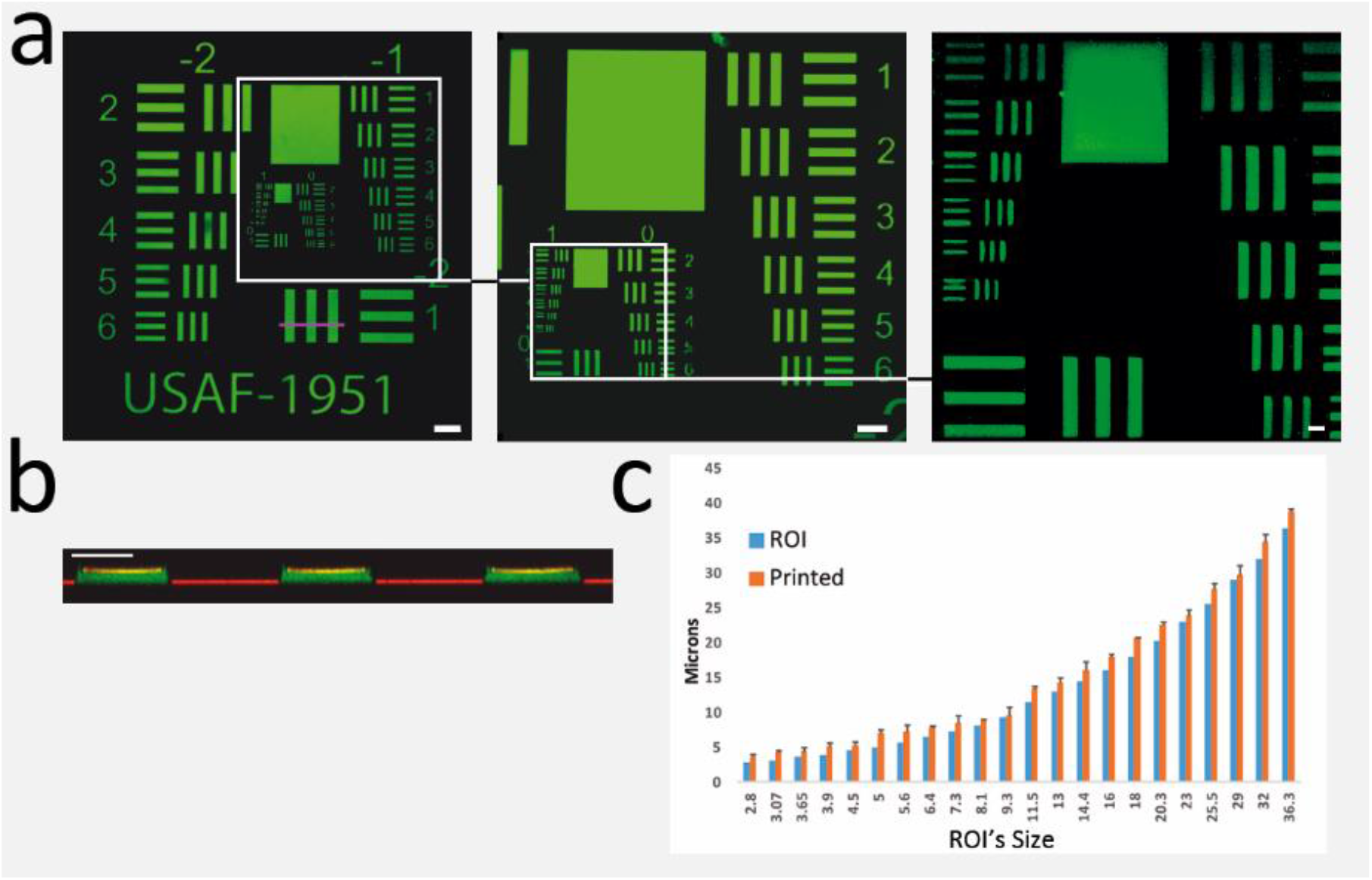
Resolution tests **a** Autofluorescent image of a printed pattern for resolution test, full print was captured with 10x magnification objective (left, scale bar 100 μm) and higher magnifications of different regions (centre, scale bar 50 μm, and right, scale bar 10 μm). **b** Z cross-section (purple line drawn on part a) showing the printing thickness, 6.5 μm. Scale bar 25 μm. **c** Quantification of all the image regions, comparing the designed ROI (blue) with printed regions (orange). Each value in the graphic corresponds to the average of three different measurements, and the error bars show the standard deviation between them.

Concerning the magnification that should be used for printing, any microscope is equipped with several objectives, each of them with a specific numerical aperture (NA). The NA of the lens affects the illumination cone. Despite of the fact that in most of the cases, a sharped focused light cone is needed (low NA), this NA variation may be used for alternative purposes. For conventional printing, we recommend using low numerical aperture objectives (5X NA 0.15, 10X NA 0.4), since the printing shape will be more uniform along with the device’s height. Additionally, this kind of objectives covers up to a 3 mm square region, speeding up the process and reaching an XY resolution up to 3 microns (Figure 3).

### Applications

Some possible uses and applications of these confocal photo-polymerized micro-devices, which we have tested in our laboratory, are described below.

#### Cell Culture

Biocompatibility of printed devices was tested with five different cell types along with this project: U2OS cell line (U-2 OS (ATCC® HTB-96™)), HeLa cell line (HeLa (ATCC® CCL-2™)), Jurkat cell line (Jurkat, Clone E6-1 (ATCC® TIB-152™) and primary lymphocytes. Cells in properly washed micro-devices have been cultured for more than 7 days with no cell viability compromise, allowing its use not only for short but for long term studies without device detaching or any resin’s pattern disturbance. Representative pictures of U2OS cell culture are shown in Figure 4a and Suppl. Material 02.

**Figure 4.**
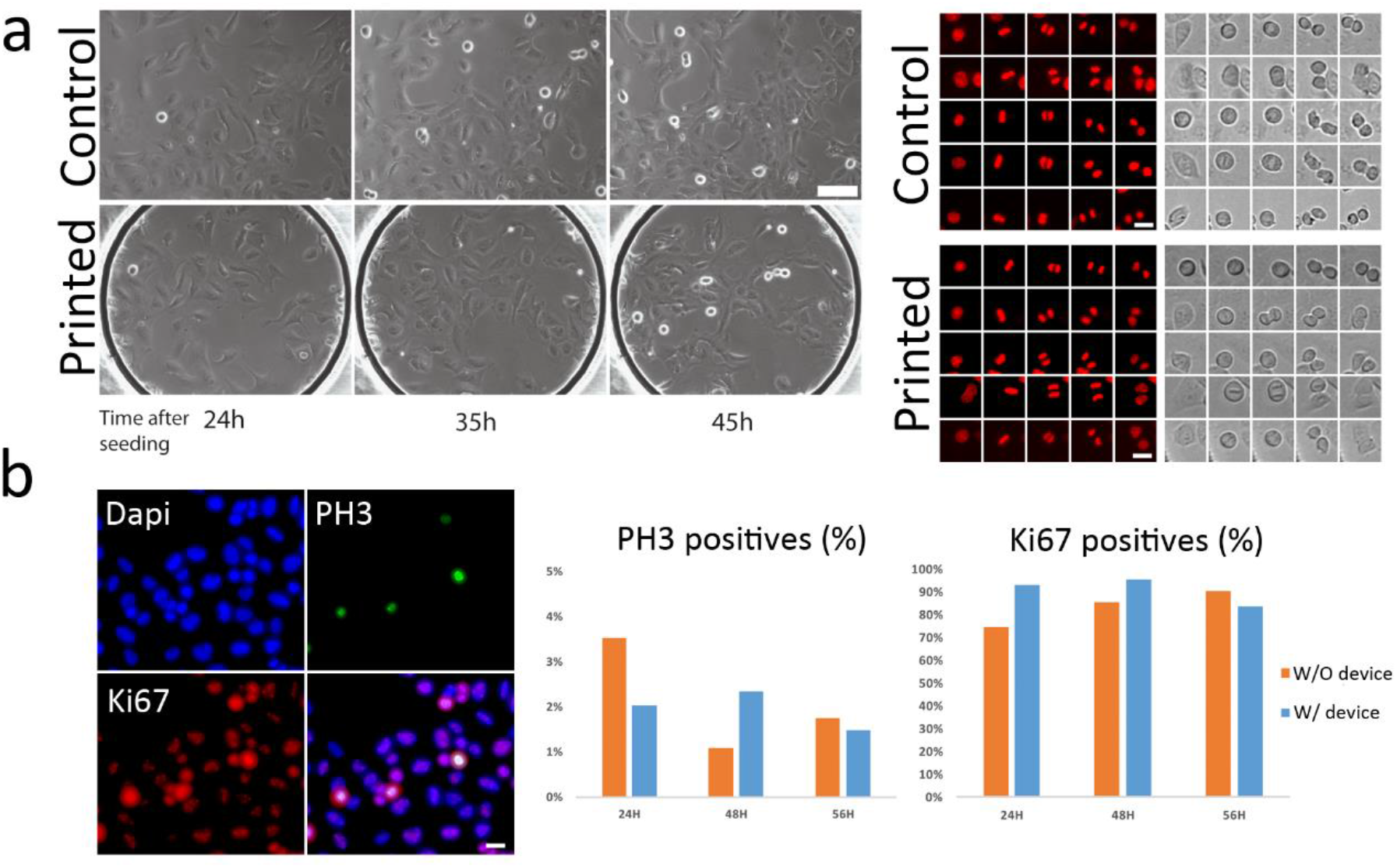
Viability **a** (left) Bright field pictures of a U2OS cell culture at different times after seeding, (right) immunofluorescence and brightfield images of mitotic events comparison between control samples and cells growing on printed devices. Scale bar left, 100μm. Scale bar right, 20 μm. **b** Representative immunofluorescence images and quantification of ki67 (proliferative cells) and Phospho-histone 3-Ser10 (mitotic cells) of U2OS cells on control and printed device. Scale bar, 25 μm.

For comparative quantification of cell viability, a study between U2OS cells cultured in printed devices vs conventional plates was performed. The immuno-staining’s of phosphorylated Histone 3 (mitotic cells) and Ki67 (proliferative) did not show significant differences. Quantification of PH3 positive cells showed an average difference of 1%, and a maximum difference of 1.5 %, between cells growing on a control coverslip without device and a coverslip with a device printed on it. On the other hand, quantification of Ki67 positive cells showed an average difference of 11.7%, and a maximum difference of 18.5 %, between cells growing on a control coverslip without device and a coverslip with a device printed on it. Further results are shown in Figure 4b.

#### Immuno-labelling

Photo-printed micro-devices show a dim auto-fluorescence in different emission wavelengths (Figure 5a). This depends on the resin used, but in general its intensity is weak, and it allows their use for bright-field applications as well as for fluorescence assays (Figure 4 and 6). Although resin auto-fluorescence can be totally bleached if needed by exposing the device to UV light for some minutes (Figure 5b), for some applications it is better to keep the dim autofluorescence, to image the micro-printed walls or pillars (Figure 6b and Suppl. Material 03). We have checked that it is also possible to specifically stain fluorescently the photo-printed devices using chemical dyes, such as CMTMR, a cell-tracing marker (Figure 9b). Although the autofluorescence from the polymerized resin is low, we have tested it with lambda scans using different laser lines to cover the whole visible emission spectrum. The experiment was performed from different regions and for three different coloured resins (black, yellow, and white) with different compositions (see materials section), reaching the conclusion that each one has different auto-fluorescence properties. In this experiments the yellow resin showed the dimmest signal, while the black resin has an emission peak in the green/orange part of the spectrum. On the other side, the white resin has the highest signal in the blue spectral region (Figure 5). So, the user can choose one or another depending on their needs.

**Figure 5.**
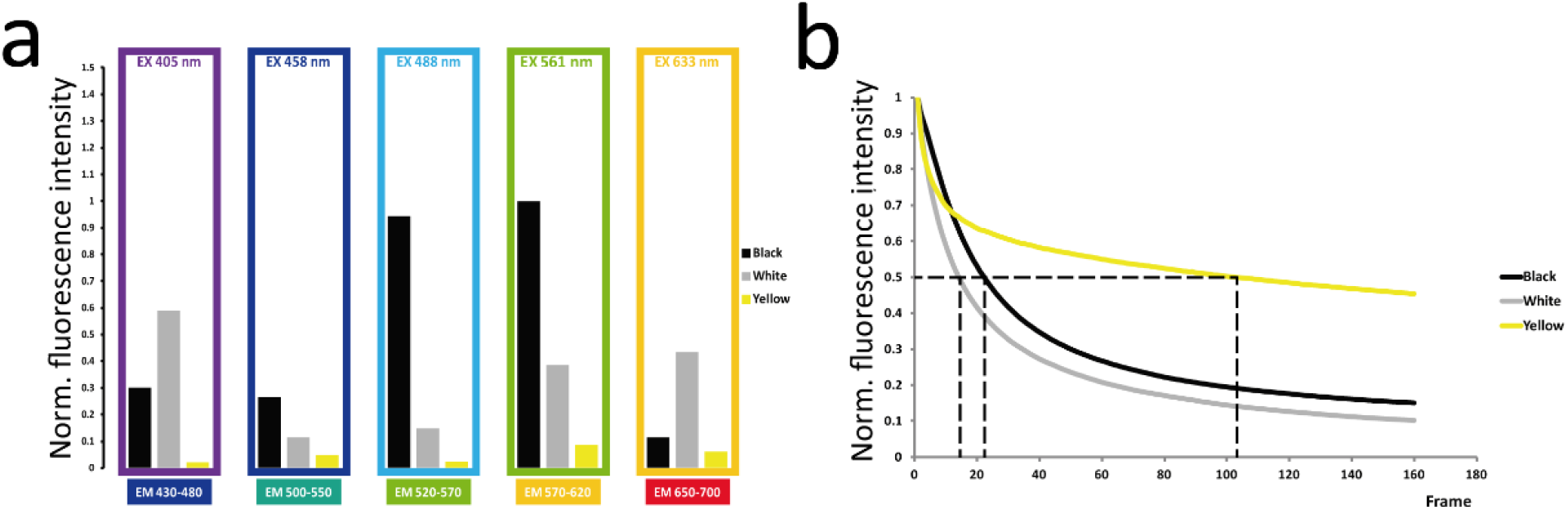
Auto-fluorescence and Bleaching **a** Auto-fluorescence levels of different resin types, normalized to the maximum level, yellow resin shown the dimmest signal while white and black were brighter in green/orange spectral regions **b** Auto-fluorescence signal decays under bleaching conditions. The yellow resin auto fluorescence needs more light to be bleached than the other two resins. Its weak autofluorescence must be considered though.

**Figure 6.**
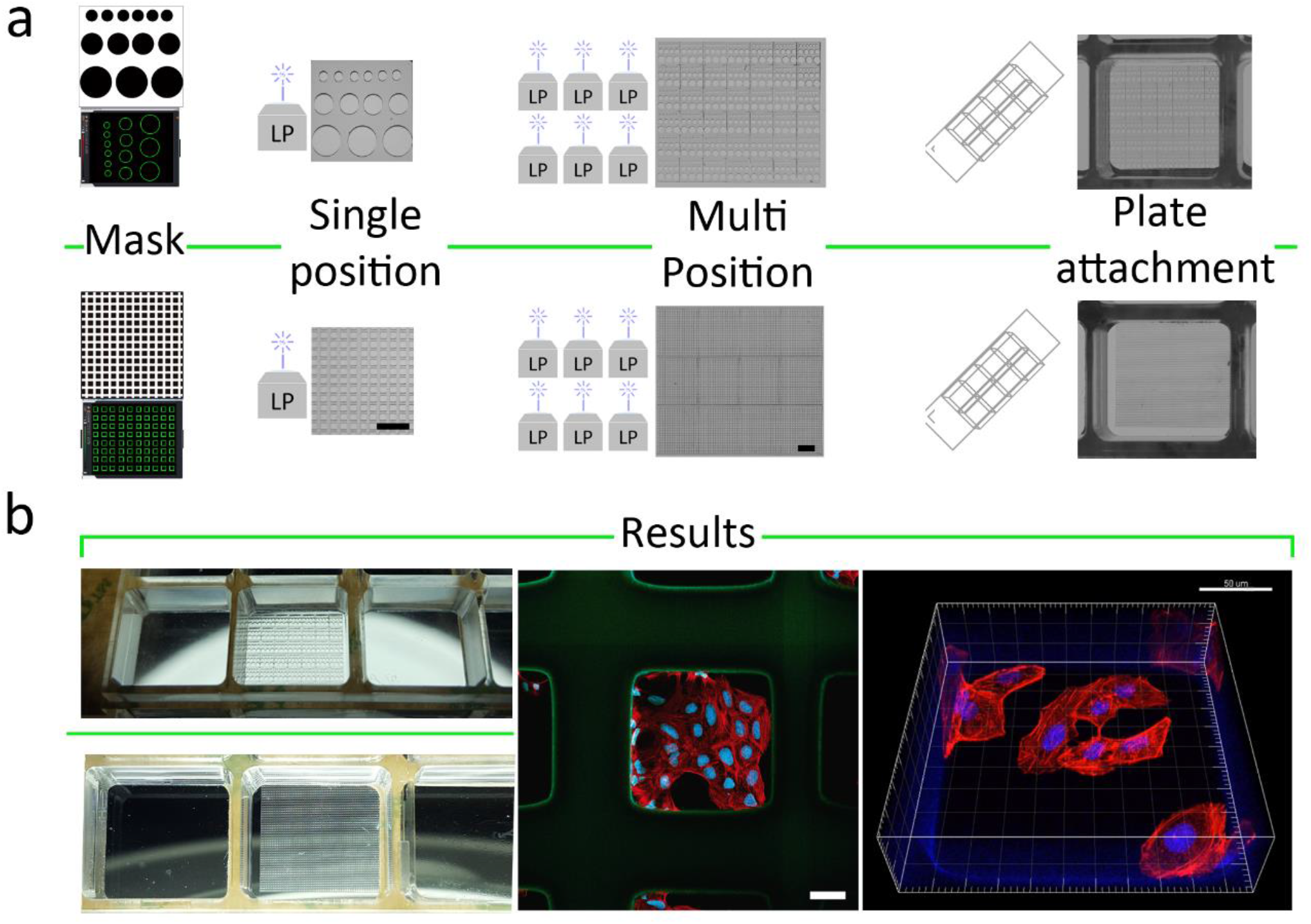
Micro-wells fabrication **a** Workflow from mask creation (left) to print (laser print, LP) in one or multiple positions (centre), until plate attachment (right). Brightfield images were captured with a magnification glass. Scale bars, from left to right, 500 um and 1 mm **b** (Left) Picture of finished results, (centre) representative picture with immunofluorescence of HeLa cells (nucleus in blue, actin in red), (right), 3D fluorescent reconstruction of one micro-well with mentioned HeLa cells inside. Scale bars are 50 um.

#### Well polymerization

The photo-printing process can be done on glass surfaces, but it can also be performed using many other formats, such as commercially available multi-well plates or channelled plates, expanding their possibilities to a wide range of different assays (Figure 6 and 7).

**Figure 7.**
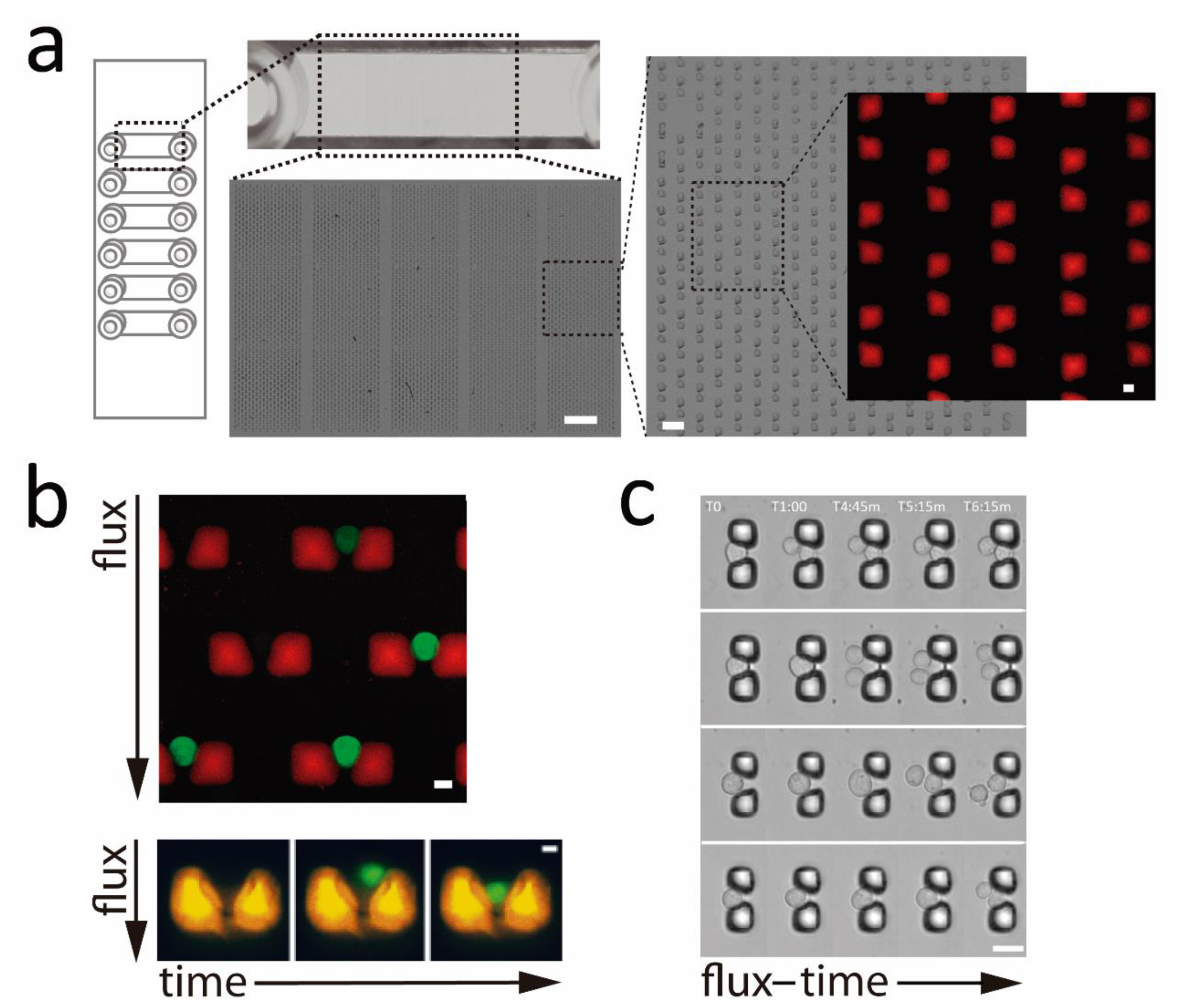
Single cell sorting **a** Detail of a single-cell-traps device printed inside a commercially available slide with channels. Medium containing cells flows inside the channel and the traps retain cells according to their size. Scale bars are 500 μm, 60 μm, and 8 μm from left to right. **b** Fluorescent images showing the results after cells flow, (up) traps (autofluorescence in red) with U2OS cells (CFSE staining fluorescence in green), (down) time lapse of how a cell is trapped. Scale bar 8 μm. **c** Time lapse of Jurkat cells divisions on single cell traps, Scale Bar 25 μm.

Additionally, a printed micro-device can be attached to commercial sticky plates (see materials section). This makes the printing, handling, and washing processes much easier, and the final device would be included inside commercial well plates just as in a common experiment with cells (Figure 6).

#### Single cell sorting

It is possible to print devices that are suitable for single cell sorting^1-4^ (Figure 7a). This can be useful to separate two cell populations with differentiated sizes (Suppl. Material 04). For this application, it may be useful to print the device on a glass coverslip, following the protocol proposed above, and then attach the glass coverslip with the device to a commercial sticky channel plate.

Live cell imaging experiments of cell traps with Jurkat lymphocytes (Jurkat, Clone E6-1 (ATCC® TIB-152™)) were also performed, with the aim of keeping them static inside traps so consecutive cell mitosis studies of cells in suspension could be tracked, making possible to measure cell cycle and mitosis timing and mitotic aberrancies (Figure 7b and 7c).

#### Multi-pattern devices

Once the field of view is photo-polymerized with the required design, the same pattern can be printed in an adjacent field. This step can be done manually, but it is much easier and precise if the microscope has a motorized stage. In this latter case, a mosaic of the same repeated pattern can be defined to automatically enlarge the polymerized region (Figure 6). What is more, you can divide a bigger pattern in several microscope fields of view, and automatically print the whole mosaic (Figure 8a). For this purpose, more complex designs are commonly required, so we developed a macro (Suppl. Material 05) programmed for its use on ImageJ open source image analysis software^34^. The macro translates any binary image that you feed into illumination ROIs, which can then be loaded in the microscope software (Figure 8a). After some software tune-up, a mosaic of photo-polymerization areas that stitch together will have been created in order to produce a complex design (Figure 8).

**Figure 8.**
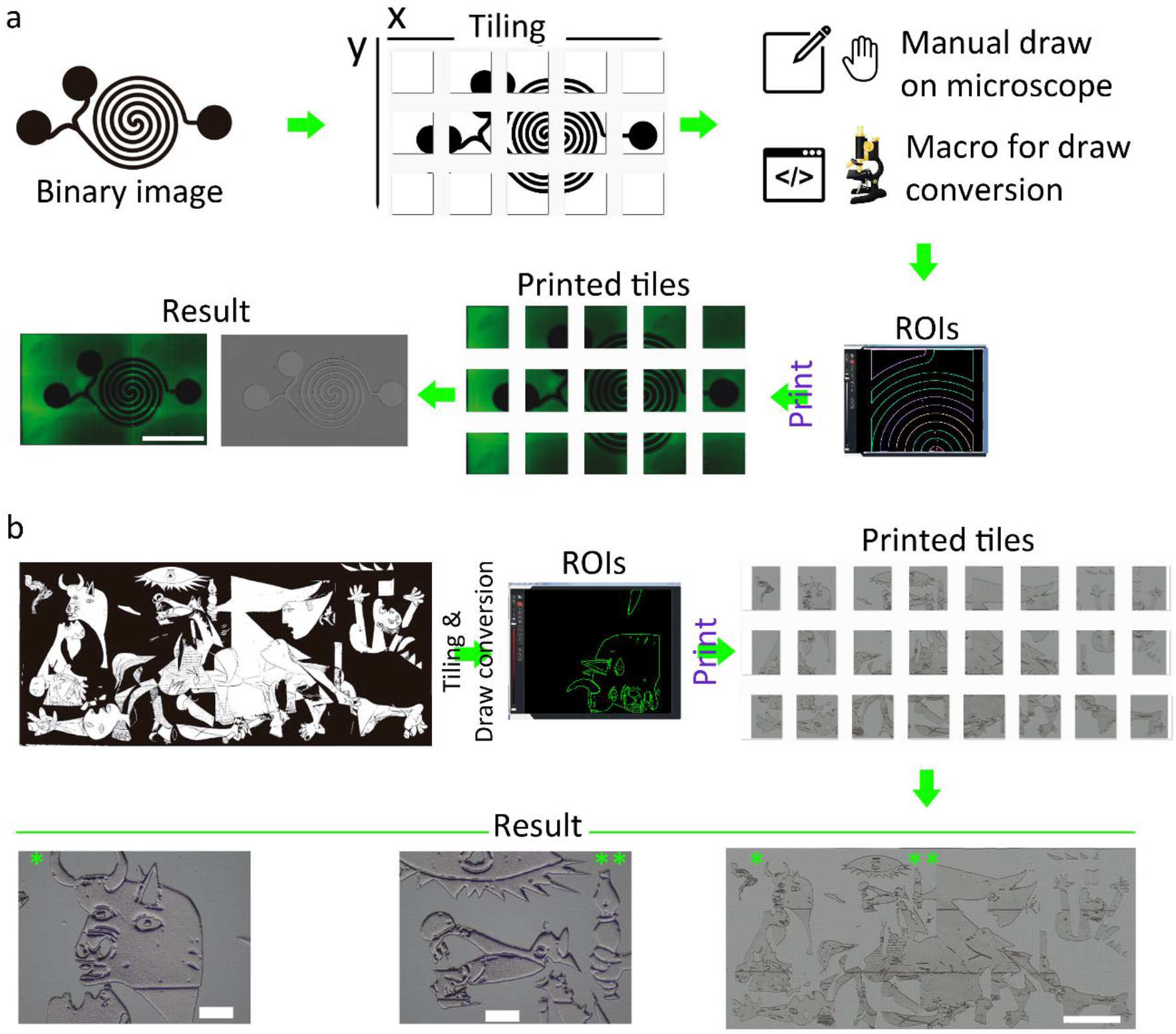
Multi-pattern print & flexibility **a** Workflow example for complex multipattern devices fabrication. The image must be separated in tiles and then each tile is translated into illumination ROIs. Tiles size depends on the scale desired and the objective that will be used on the microscope. This process can be done manually but the use of a customized macro (example included in supplementary material) simplify this task. Green result image and printed tile images are autofluorescence, and gray result image is brightfield. Scale bar, 5 mm. **b** Example of flexibility. By following the instructions above, almost any design is possible. In this case, a brightfield image of a 2 cm-length printing of the Picasso’s Guernika painting, is shown. Scale bars, 2.5 mm for the whole printing, and 500 μm for detailed pictures.

This allows to generate devices for any microfluidic assay with highly complex designs (Figure 8a and Suppl. Material 01 and 06). In this kind of approach, the device must be properly sealed, as previously described above. The complexity of a micro-device printed this way can range from very simple designs to really complex ones (Figure 8b).

#### Multiphoton-printed scaffold for 3D culture

While the method described here is focused on the use of regular confocal microscopes, a similar approach can be followed by using multiphoton microscopes (Figure 9). In this regard, the advances in photo-polymerization using multiphoton microscopes are especially interesting^10,13,28,29,35^, giving the chance of printing 3D shapes in the Z dimension. However, this kind of equipment is not so easy to find, it is much more complex to use and, in our experience, it doesn’t grant an improvement in terms of XY resolution. In our experience, we found that multiphoton photo printing requires specifically designed resins^28,35^ and high-power laser, thus limiting the device size and increasing the fabrication time. Nevertheless, confocal and multiphoton photo printing can be combined if needed.

**Figure 9.**
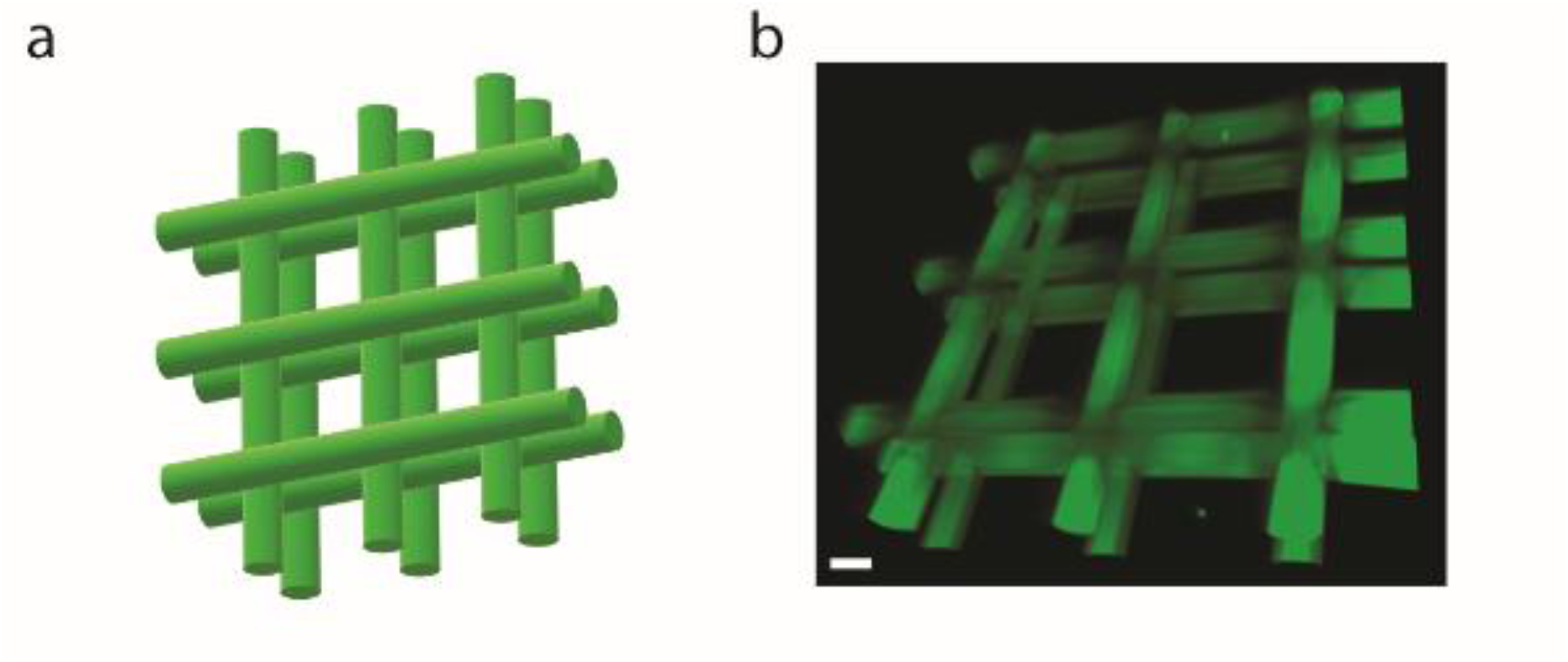
Multiphoton-printing **a** 3D schematic representation of a scaffold structure intended for 3D cell culture. **b** Final 3D result of how the scaffold looks like after printing, washing and staining with CMTMR dye. Scale bar is 20 μm.

#### Mold for conventional micro-devices fabrication

Finally, while this method is intended to create micro-devices ready to use without the need of generating a previous mold, another possibility is to generate a negative relief that can be used later for conventional PDMS devices fabrication^6,14,15,17-19^.

## Discussion

This novel method presented here allows fabricating micro-devices for cell-based assays to any laboratory without any dedicated equipment, simply by using a laser scanning confocal microscope and a few and inexpensive reagents. Obviously, this protocol relies on the use of an expensive machine such as a confocal microscope, but this is common equipment nowadays in most research institutes and universities and is usually available in specialized core facilities. The pursued workflow would be to design illumination ROIs by using conventional drawing tools or an offline version of the microscope software and then booking a few minutes in the nearest available confocal microscope for the photo-polymerization process. In addition, the flexibility of this method is a key point, since several prototypes can be created for testing within minutes, to get the definitive design. Moreover, it also can be used to create master molds suitable for conventional production.

With this approach, single-cell traps, flux experiments and long-term cell culture can be achieved effortlessly, without the need for special conditions for storage. Immunofluorescence assays can also be performed using these micro-devices, allowing the possibility of multiple staining experiments in a reduced space, which can be useful for studies with a limited number of cells. Considering the short time needed for the microscope printing, and the cost of the reagents, the result is an extremely affordable and fast protocol that may finally bring micro-devices fabrication closer to many cell biology laboratories, regardless of their familiarity with this technology. By applying this combination of photosensitive resins and microscopy, more applications will appear in the near future, such as for example to introduce landmarks in correlative light-electron microscopy (CLEM), or microscopic particles generation.

## Materials & Methods

### Materials used for printing

In this project, three different standard 3D printing resins were tested, each of them with different properties and different printing efficiency. One of them was colorized with a black pigment (made solid ms resin v2 (msr1lobv2) black), the second one was colorized with a white pigment (made solid white vorex tough resin (mstufltw0), and the last one with a yellow pigment (made solid cast solid resin (mscs)). White resin was the first to trigger polymerization, followed by the other two. This could be something to consider in case of having a weak illumination source. For the purpose of this publication, the black resin was mainly used.

In order to improve the adherence of the polymerized resin to the printing surface, 3-(Trimethoxysilyl)propyl methacrylate (TMSPMA) (Sigma Aldrich 440159) was used, as previously described.

To prepare a mix of resin, in order to improve the spreading over the printing surface, diethylether was used as a solvent (Sigma Aldrich 673811)

In all printing processes, TCS SP5 X AOBS and a TCS SP5 AOBS (Leica microsystems), both equipped with AOBS and AOTF to allow ROI-specific control of illumination, were used. The lasers used were a 405 SSL, an argon (458-514), a 561 SSL and a 633 HeNe. Additionally, the TCS SP5 AOBS confocal had a multiphoton laser. For the polymerization process, 405 nm laser illumination, and 5x HCX PLAN FLUOTAR NA 0.15 and 10x HCX PLAN APO CS NA 0.4 objectives, were used. To determine the level of polymerization, auto fluorescence was collected between 470 and 545, using 405 and 458 combined for the excitation. Since solid resin increases its auto fluorescence, the best time to stop the illumination can be determined (see suppl. Material 01). Prior to the polymerization, the focus plane must be chosen (i.e. the interface between resin and glass). For this purpose, laser reflection using a 633 nm laser-line was collected, placing the detector to collect 633 nm excitation and setting AOBS to let reflection pass at that wavelength.

After the printing process, washing of the printed device is needed, as detailed in results section. A final step of PBS washing may be required, if biocompatibility is needed.

### Cell culture

U2OS (U-2 OS (ATCC® HTB-96™)) cells were cultured in DMEM medium (31966-021 (Thermo Fisher)) supplemented with 10% fetal bovine serum (FBS), 100 units/ml penicillin, 0.1 mg/ml streptomycin, 1% glutamine and 1% sodium pyruvate. Cells were maintained at 37ºC in a humidified atmosphere of 5% CO2.

### Immunofluorescence and fluorescent staining

For the fluorescence imaging of the nucleus, sirDNA probe (Spirochrome) was used at a final concentration of 3 μM. Cells were incubated with the probe during 1 hour prior to start imaging and was also left in the cell culture medium along the duration of the time lapse. Fluorescence in the far-red zone of the spectrum was collected using a 20x HC PLAN FLUOTAR NA 0.5 objective and Y5 filter (EX BP 620/60, EM BP 700/75) on a DMI6000B widefield microscope (Leica Microsystems).

Fluorescent staining was made to quantify viability of U2OS cell line growing on a printed device versus coverslip glass. Cells were fixed at different time points, using 4% Paraformaldehyde (PFA) (Thermo Fisher 28906). After fixation of all time points, cells were permeabilized using 0.5 % Triton in PBS. Cells were blocked with TNB (Boehringer 1096176), and incubated 1h at 37ºC with primary antibodies against PH3-Ser10 (Merckmillipore, 05-806) and Ki67 (Abcam, ab16667), both at a final concentration of 1/500. After washing, incubation 30’ with secondary antibodies and DAPI (Termofisher, D1306) was performed: an Alexa 488 anti-mouse antibody (Thermo Fisher A-21202) was used to stain PH3-Ser 10, while an Alexa 555 anti-rabbit antibody (Thermo Fisher A-32794) was used to stain Ki67. DAPI (Termofisher, D1306) was used at a final concentration of 5 μg/ml. After washing, cells were left in PBS and imaged for quantification, using a widefield fluorescence microscope (Leica DMI6000B) with a 20x NA 0.5 objective and filters A4 (EX BP 360/40, EM BP 470/40) for DAPI, L5 (EX BP 480/40, EM BP 527/30) for Alexa 488 and N2.1 (EX BP 515-560, EM LP 590) for Alexa 555. Quantification of positive PH3-Ser10 and positive Ki67 cells over total number of cells for all time points was done using Definiens software (Definens).

Staining for DAPI and Phalloidin is shown for HeLa cell line. Cells were cultured with DMEM medium supplemented with 10% fetal bovine serum (FBS), 100 units/ml penicillin, 0.1 mg/ml streptomycin, 1% glutamine and 1% sodium pyruvate. Cells were maintained at 37ºC in a humidified atmosphere of 5% CO2. Cells were fixed using 4% PFA, and incubated with DAPI (Termofisher, D1306) at a final concentration of 5 μg/ml, and with Phalloidin-568 (Thermo Fisher A-12380). Pictures were captured with a Leica TCS SP5 X AOBS confocal (Leica Microsystems). Device auto fluorescence and DAPI fluorescence was captured using a 405 nm laser and collecting light between 413 and 440 nm; phalloidin fluorescence was captured using a 561 nm laser and collecting emission between 569 and 669 nm. A stack was captured using a 40x HCX PLAN APO CS NA 1.25 objective with a zoom factor of 2, and then a 3D reconstruction was made using Bitplane Imaris 7.31.1 software (Biplane).

### Resolution performance quantification

To test the resolution limit and performance of our method, a regular image commonly used to test scanners’ resolution was used. First, the image was binarized, converted to ROIs and those ROIs were printed with resin, using a 10x NA 0.4 objective. Quantification was done by, using ImageJ, tracing lines perpendicularly to the printed bars, and then the fluorescence profiles of those lines were plotted. This allowed us to measure bars’ thickness, to finally compare those distances to the original distances coming from the ROIs we loaded in the microscope software.

### Multi-position printing

For the automation of printing a multi-position device, the additional LAS AF application, named Leica Matrix Screener application (Leica Microsystems), can be very useful, and it can be added to almost any Leica SP5 confocal microscope. This allows to create and save templates with several distances and printing configurations. In any case, the result is the same if every field is positioned and printed manually, yet more laborious to do.

For the printing of complex designs that cannot be printed in just one field of view, a ready-to-use macro (Suppl. Material 05) for ImageJ^34^ has been designed. This translates any binary image to ROI files that can be easily loaded. This kind of file has been tested on a TCS-SP5 and on a more modern Stellaris and SP8 (Leica microsystems). A simple macro modification may be required if loading ROIs in a different microscope brand is needed. Macro use is described in supplementary material 05.

### Auto fluorescence and bleaching quantification

For the auto fluorescence quantification, LAS AF software was used, both for lambda scans captures and for intensity measurements. Lambda scans were done using a 20x HCX PLAN APO CS NA 0.7 objective with a zoom factor of 4.

Bleaching quantification was done with 405 nm laser on a region of 193 μm (20x objective with a zoom factor of 4) over printed and cleaned resin, fluorescence emission was collected between 430 and 540 nm.

### Single-cell sorting

To get these cell traps that are able to sort cells according to their size, the design was printed on a standard rectangular coverslip and, after washing and sterilizing steps, a commercially available Ibidi sticky plate with channels (sticky-Slide VI 0.4, Cat.No: 80608) was attached. Then, tubes from OB1 air-pressure system (Elveflow) were connected to the plate, creating a controlled flux. Next, the system was loaded with PBS, then with culture medium, and finally with culture medium containing U2OS (U-2 OS (ATCC® HTB-96™))/Jurkat (Jurkat, Clone E6-1 (ATCC® TIB-152™)) cells. Tracking of how cells were being trapped was performed by live-image acquisition in a widefield fluorescent microscope (Leica DMI6000 B), using a low exposure time to have a real-time image. To improve detection, cells were stained with CFSE (Thermo Fisher C1157), a green cell marker. High resolution images from captured cells were acquired with the confocal microscope already mentioned before, collecting autofluorescence from the traps using 405 nm excitation laser and between 430 and 493 nm for emission; emitted fluorescence between 500 and 540 nm, coming from CFSE-stained cells, was collected, using a 488 nm laser for excitation.

Additionally, polyvinyl alcohol (Sigma Aldrich 1413521000) could be used to prevent cells from getting cells stuck to plate surface. To do this, once the cell-trap device is printed, properly washed, sterilized and mounted on a sticky plate, the channelled plate was incubated with polyvinyl alcohol for 2,5 hours. After that, it was washed several times with PBS and proceeded as explained above.

### Flux assays

After sealing of a device designed for flux assay, luers couplers can be plugged to the entrance and exit of the device, and using OB1 air-pressure system (elveflow), it is possible to load liquids and cells at a controlled flux rate. As explained before, live-image acquisition at the same time that medium with CFSE-stained cells was flowing was performed using a Leica DMI6000 B microscope. A time lapse video of this experiment is shown in Suppl. Material 06.

### Multiphoton printing

To print several layers to create a 3D structure, multiphoton printing (MaiTay Laser Spectraphysics) with a 10x objective was used, since only a thin layer polymerizes during illumination. This allows to move the Z position and load different ROIs in each layer, resulting in a more complex 3D design that is not possible to be created with conventional lasers, since the cone of light polymerizes from top to bottom of the resin layer.

After the printing process, the 3D scaffold was stained with CMTMR for 2 minutes (C2927, Thermo Fisher) and washed with PBS. The result was a brighter fluorescence coming from the device, allowing to have a better signal for imaging. A stack of this was captured with a confocal microscope, and a 3D reconstruction was generated using Bitplane Imaris 7.31.1 software (Bitplane).

## Supporting information

Supplementary 5 Guide

Supplementary 5 ImageJ Macro

## Acknowledgments

We would like to thank to Tatiana Grazioso, Laura Pena, José González, Aleida Pujol, Cristina Tejedo, Daniel González and CNIO’s Flow Cytometry Unit for kindly contributions in all of our in vitro biocompatibility assays. We are also very grateful to Fernando Peláez and Arrate Muñoz, for their invaluable help with the review of this manuscript. Last but not least, we also want to thank to David Martín, from Leica’s microscopy division, for lending us imaging instrumentation, and Alessia del Piano, for contribution with some good ideas at the beginning of this project. To all of you, thank you for your inestimable help with making this possible.

