## Supplementary 5 Guide for "Laser Scanning Confocal printing, a new simple method for micro-device fabrication (LSCprint method)"

### RoiLoader User Guide

Developed by Javier Ortega

In the Confocal Microscopy Unit of the CNIO

Last Updated: 16/12/2020

### INTRODUCTION

Laser scanning confocal microscopes take advantage of a set of mirrors to move a laser across the specimen. The path of this laser in the sample can be modified by drawing ROIs and imposing to the scanning systems to cover just the area inside (or outside) the ROI.

In the particular case of Leica's confocal microscopes, the software that controls the system, LAS AF, has very limited drawing tools, confining the ROI drawing to very basic shapes.

The RoiLoader plugin is able to translate a binary image, or image stack, opened in Fiji into a .roi file that can be understood by the microscope. Thanks to it, it is possible to work with more complex ROIs to, for example, build microfluidic chips using the confocal microscope.

### BASIC CONCEPTS

To understand the macro, it is necessary to know some basic concepts:

- Design: Binary image opened in Fiji.
- Tile: Area of the sample that can be imaged with a certain objective at once in the microscope.
- Tiling process: Process that consists in the division of the design into subimages, each one of them corresponding with the size of a single tile.
- ROI: Region of Interest in an image, it is referred to the particles detected by the *Analyze Prticles* function in Fiji and to the regions drawn in LAS AF.
- Group of ROIs: Set of ROIs that can be handled as one in the microscope.

### OVERVIEW OF THE MACRO

The macro will first perform the tiling process, and for each tile it will detect all the rois with the Analyze Particles function. These rois are going to be grouped and then, all the information will be stored in a .roi file.

At the end, a folder will contain:

- A smaller copy of the design.
- A log .txt that will include the settings selected to obtain the rois and any extra comments made by the user.
- A folder for each slide in the image stack. Inside them, a .roi file for every tile. Each file will be named as follow:
  'Name + z position+ x position + y position' where the positions correspond to the original design. The tiles that correspond just to a white square will be named with 'Empty' at the end of the standard name to make easier their use. In the case of a single, non-stack, image, there will be no folder for the z.

#### Input design

The input design has to be a binary image, allowing image stacks, and the rois detected will correspond to the white (255 pixel value) areas in it. In order to assure that white and 255 pixel value correspond to the same areas in the image, the LUT must not be inverted. The program will check if the image is binary and has a non-inverting LUT, if the design does not fulfill these requirements, the plugin will stop and should be executed again when the image is corrected.

- Image is not binary.

If the image is not binary, the following message will appear:

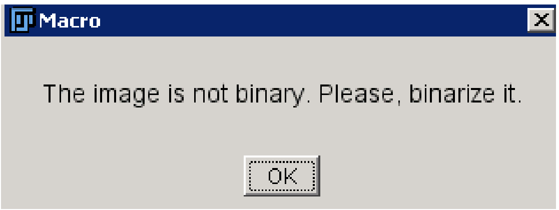

To fix it: Process>Binary>Make Binary to perform an automatic binarization or Image>Adjust>Threshold to set a threshold to the binarization.

Be aware that during the binarization process the LUT may be inverted.

- LUT is inverted.

If the LUT is inverted, the following message will appear:

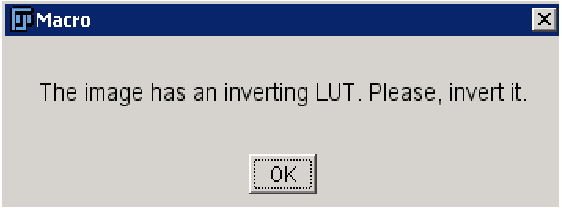

To fix it: Image>Lookup Tables>Invert LUT.

Please note that it will be necessary to invert the image in order to keep the same white areas as before inverting the LUT.

#### Dialog Box

When the macro starts, it shows a dialog box where the user must enter the needed information, it is divided in three blocks. The first, asks for the directory and the name of the file, it will have as a default parameter the directory and name of the loaded image, with the possibility of displaying a window to browse for the directory after clicking OK. The final name of the file will be the selected name followed by the position of the tile with respect to the full design (Name+ X position+ Y position). The second one, asks for the scale of the design, where the user must enter a known distance corresponding to a known number of pixels in order to calibrate the design. Finally, the information about the tile size and the overlap is required. There is an extra space in the box that allows any extra comments that will be written in the final log file.

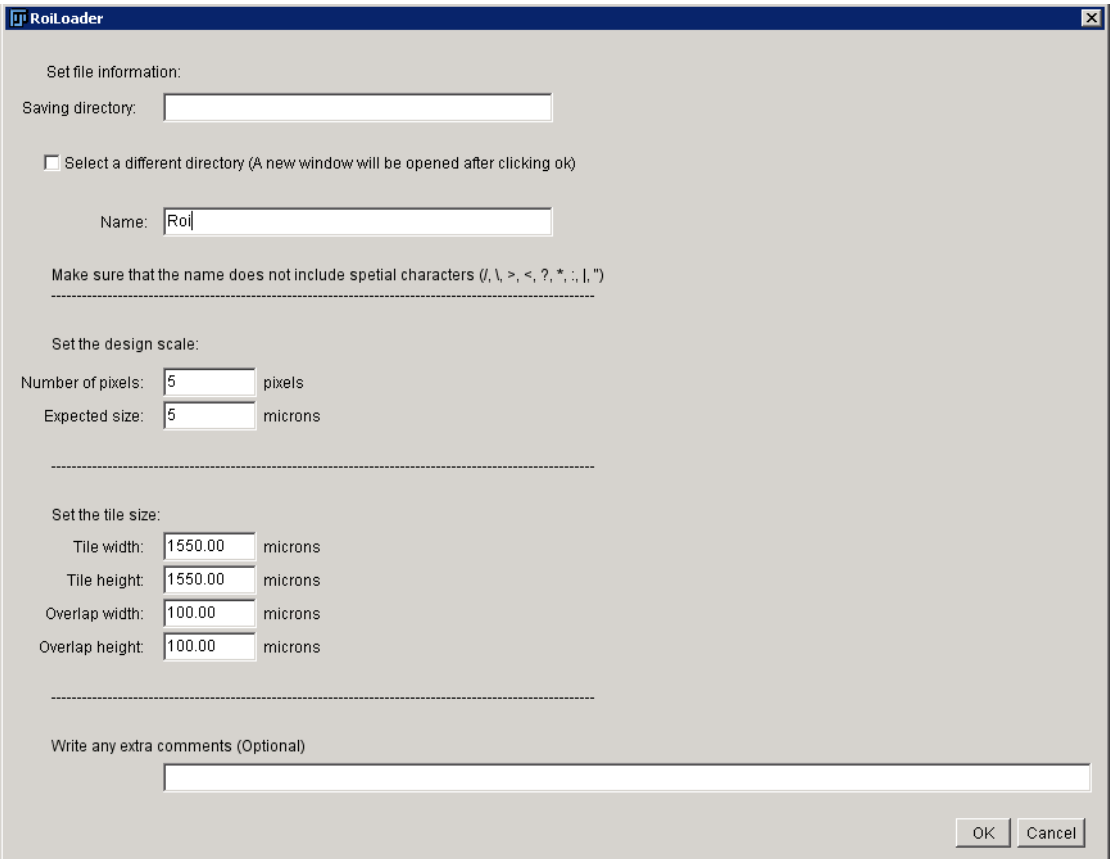

The numeric values entered should be positive, and the overlap can also be 0. If these values are wrong, it will ask for the inputs again:

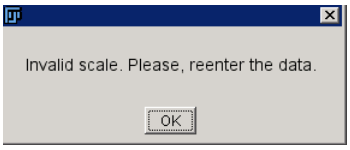

If another directory is wanted, after clicking OK, it will appear the next window, where the user can select the desired directory to save the results:

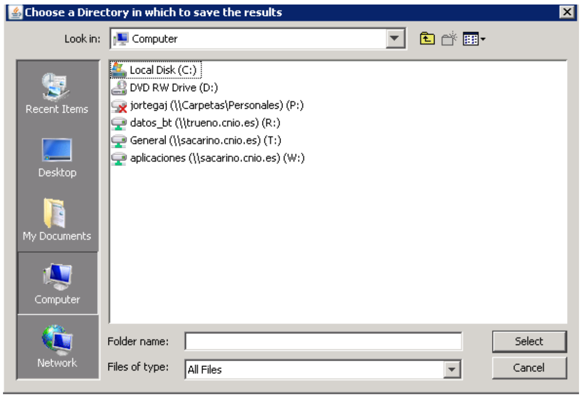

#### Scale

To enter the scale, the user has to give a known pixel-microns relation. For example:

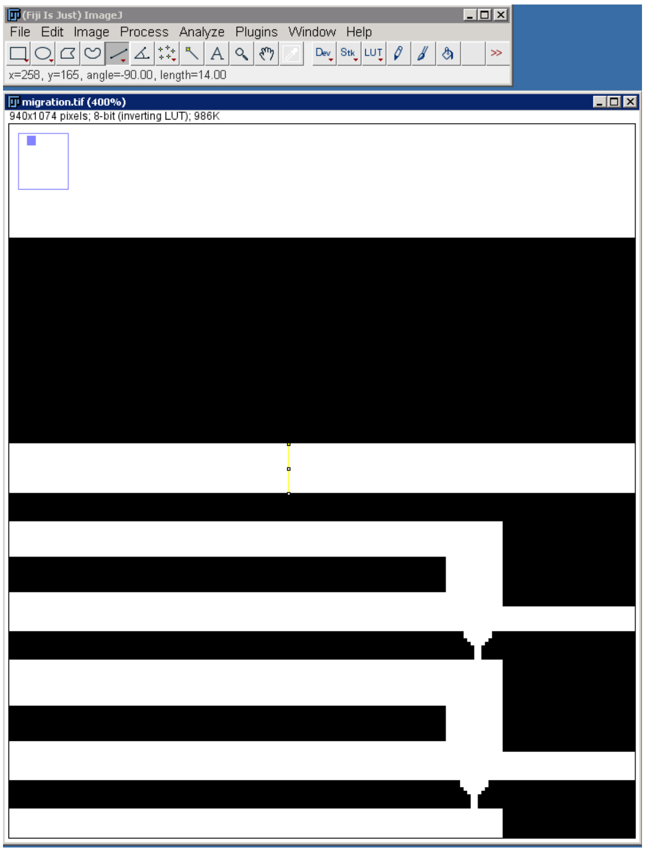

It is possible to check that the channel has 14 pixels, and we know that the channel must has 30 microns width, then in the dialog box it must be entered:

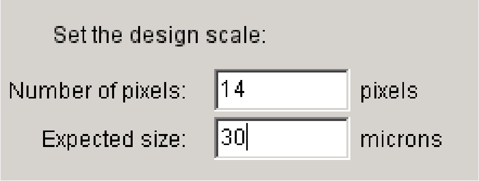

#### Tile size

The tile size in microns is expected to match the tile size of the microscope using the same objective as the one to be used when working with the generated roi file.

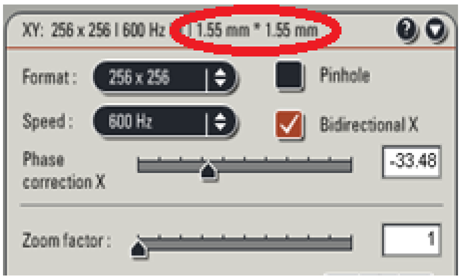

In this example, the correct setting will be:

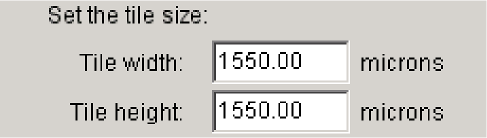

Notice that the input is asked in microns, while the tile size in LAS AF can be in millimeters.

#### Overlap

The overlap must be set in order to allow a proper tiling process. Each microscope has its own inter-tile distance, so this parameter can be modified in the dialog box or it can be set to 0 and modified in the microscope software.

After clicking Ok, all the parameters will be verified and, if they are correct, the macro will create the .roi files.

#### Final result

Once the macro has ended, a message will appear showing the path where the results will be found.

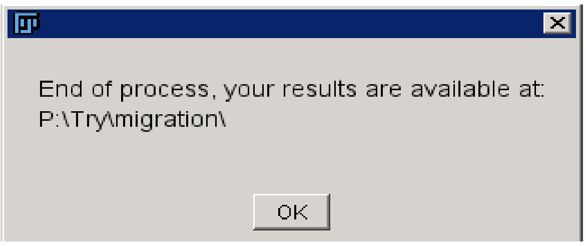

In the selected path, we will find the folder with the results:

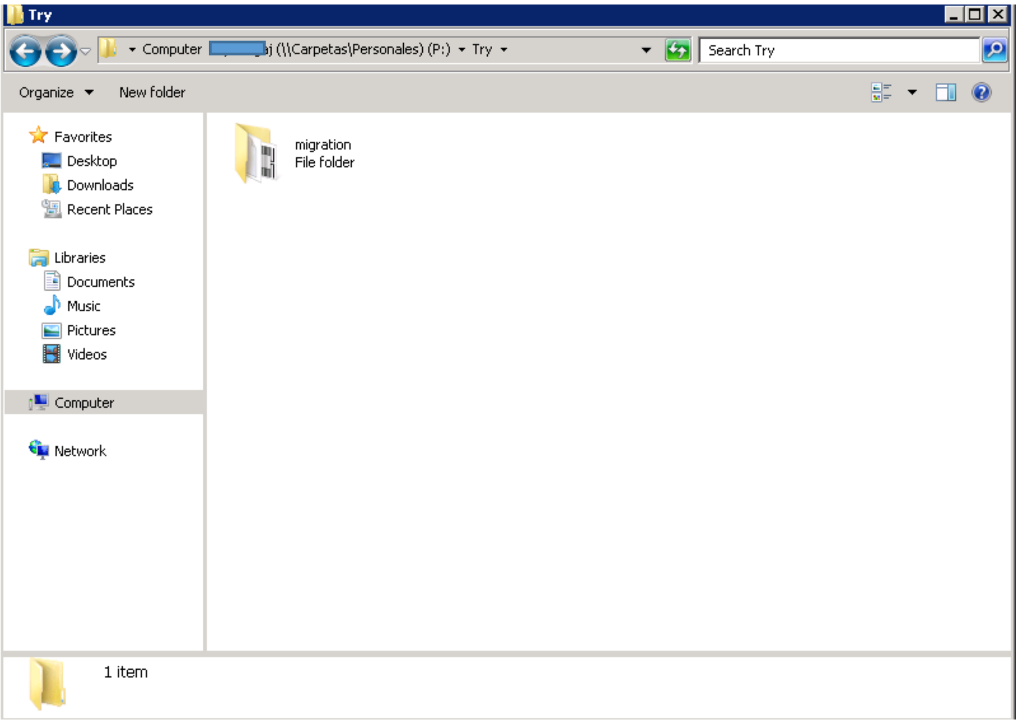

Opening the folder it can be found four .roi files named with the experiment name followed by the position of the tile in the full design, the log file and a copy of the design:

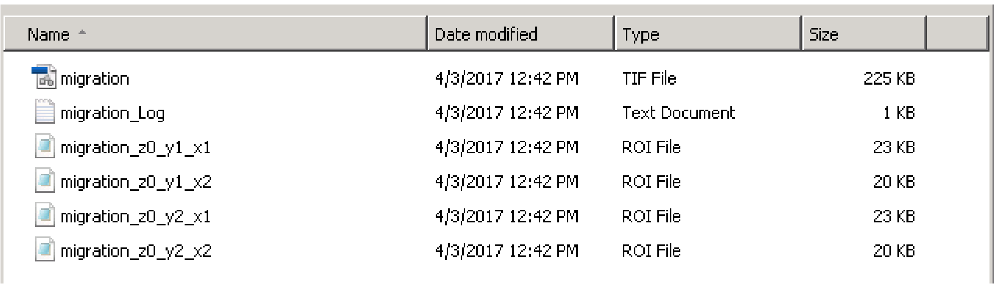

| Name | Date modified | Type | Size |
|---|---|---|---|
| migration | 4/3/2017 12:42 PM | TIF File | 225 KB |
| migration_Log | 4/3/2017 12:42 PM | Text Document | 1 KB |
| migration_z0_y1_x1 | 4/3/2017 12:42 PM | ROI File | 23 KB |
| migration_z0_y1_x2 | 4/3/2017 12:42 PM | ROI File | 20 KB |
| migration_z0_y2_x1 | 4/3/2017 12:42 PM | ROI File | 23 KB |
| migration_z0_y2_x2 | 4/3/2017 12:42 PM | ROI File | 20 KB |

The Log file has the following information:

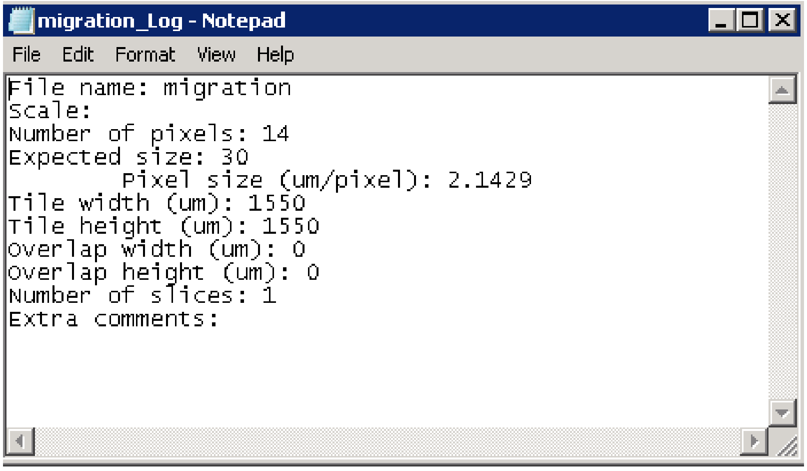

In the case of a stack image, the results will be found as follows:

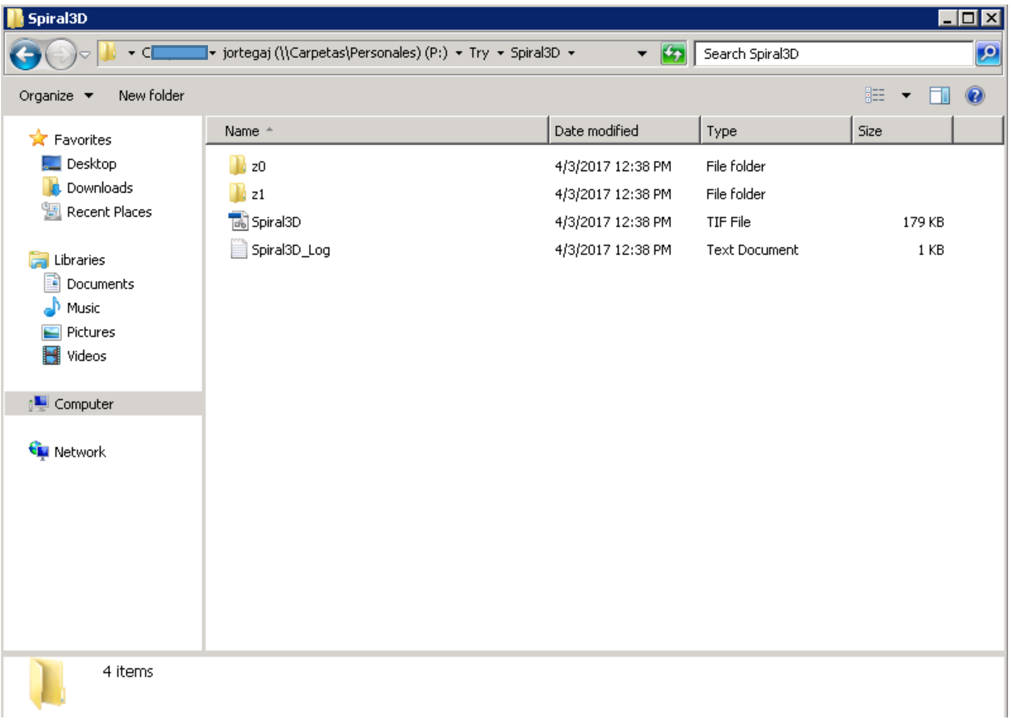

And inside each folder:

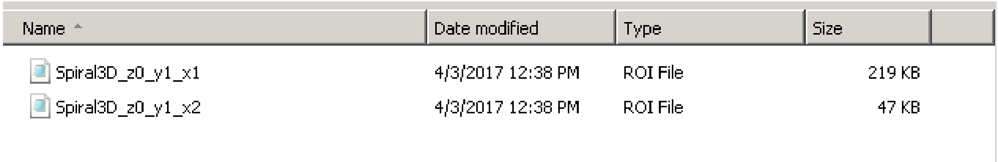

The Log file also shows the number of slices:

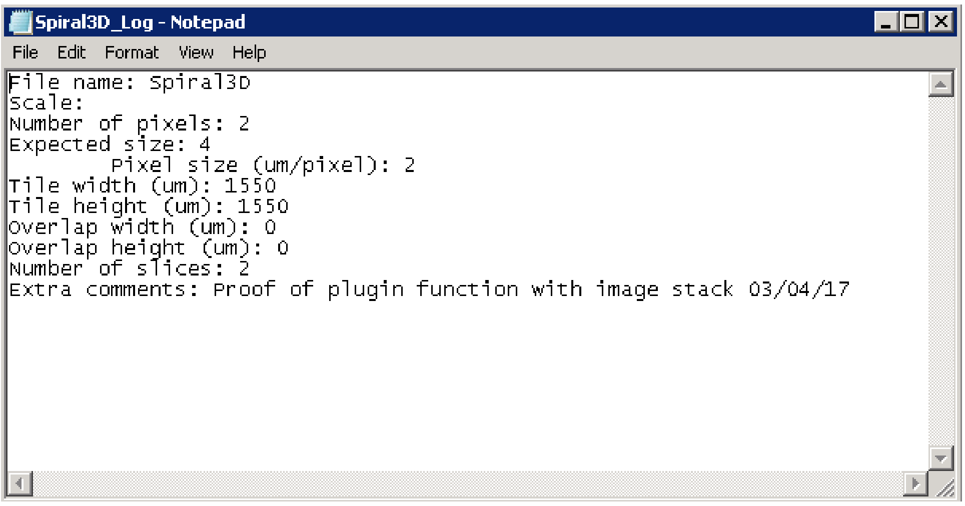

#### Problems and known issues

- Maximum number of ROIs.

Due to code efficiency optimization, the number of rois per tile that can be detected is 10000. If more rois are found, it will appear the error:

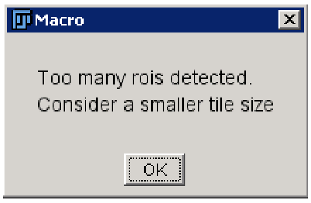

This error can be solved by two different approaches. First is considering a smaller tile size, so that there are fewer rois on it. The second one allows the detection of all the rois in the tile without changing the size, but involves modifying the code.

**Code modification:**

A key step in the code is to group the rois to ease their handle in the microscope. To group the rois, they must be firstly sorted by the size of their bounding rectangle. The macro consider the names of the rois as strings, so they must be sorted alphabetically. At line 164 of the code, the ROIs have been renamed according to its bounding rectangle area, the biggest will be called 0, the next one 1… The problem arises when names with different number of digits are included, as the sorting will be done: 1,

10, 11, 12…2, 3, 4… To avoid it, their name must be changed adding 0s in front of the number with the function:

**roiManager("rename", name)**

Which renames the selected ROI so that they are sorted: 01, 02, 03…10, 11, 12… This change is done by the code:

```
roiManager("select", reversedRankedBoundingRectArea[i]);
if (i<10)
{
    roiManager("rename", "000"+i); //the 0s are added in order to make the sort, as the names are considered as strings.
}
else if (i<100)
{
    roiManager("rename", "00"+i);
}
else if (i<1000)
{
    roiManager("rename", "0"+i);
}
else if (i<10000)
{
    roiManager("rename", i);
}
else
{
    selectWindow("tile");
    close();
    selectWindow("temp_image");
    close();
    selectWindow("ROI Manager");
    run("Close");
    selectWindow("Log");
    run("Close");
    selectWindow("Results");
    run("Close");
    exit("Too many rois detected.\nConsider a smaller tile size")//If there are too many rois, the macro will be aborted.
    //Add more else if to consider a larger number of rois  if needed.
}
```

As it is seen, the code just includes the change for fewer than 10000 rois.

To change it, add after the last *else if*:

else if (i<100000)

{

roiManager(“rename”, i);

}

And add one “0” to the previous rename arguments.

The code should look like (changes highlighted):

```
    for(i=0; i<roiManager("count");i++)//rename the rois according to their bounding rectangle area rank.
    {
        roiManager("select", reversedRankedBoundingRectArea[i]);
        if (i<10)
        {
            roiManager("rename", "0000"+i); //the 0s are added in order to make the sort, as the names are considered as strings.
        }
        else if (i<100)
        {
            roiManager("rename", "000"+i);
        }
        else if (i<1000)
        {
            roiManager("rename", "00"+i);
        }
        else if (i<10000)
        {
            roiManager("rename", "0"+i);
        }
        else if (i<100000)
        {
            roiManager("rename",i);
        }
        else
        {
            selectWindow("tile");
            close();
            selectWindow("temp_image");
            close();
            selectWindow("ROI Manager");
            run("Close");
            selectWindow("Log");
            run("Close");
            selectWindow("Results");
            run("Close");
            exit("Too many rois detected.\nConsider a smaller tile size")//If there are too many rois, the macro will be aborted.
            //Add more else if to consider a larger number of rois  if needed.
        }
    }
```

Now it will detect up to 100000 ROIs.

To further include more rois, perform the corresponding changes in the same way.

- The ROI does not appear in the LAS AF window

If when the ROI is loaded into the microscope it does not appear in the window, it can be a problem related with the scale or the tile size. Check if the introduced data corresponds with the tile size that is being used in the microscope.

- Java Error

It is possible that during some executions it prompts the following error message:

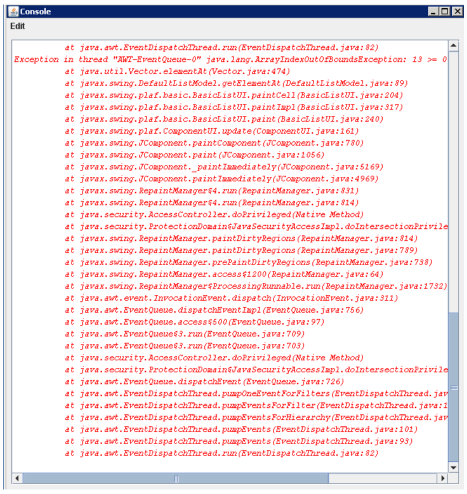

However, this error does not affect the macro function, so it can be ignored.
