## Supplementary 5 ImageJ Macro for "Laser Scanning Confocal printing, a new simple method for micro-device fabrication (LSCprint method)": Suppl. material 05_RoiLoader_.ijm

```
//The RoiLoader macro has been developed by Javier Ortega Juliá at the Confocal Microscope Unit of the CNIO using Fiji (ImageJ Version: 1.49r)
//This macro will transform a microfluidic chip design in one or more slices (or any other binary image) opened in Fiji, into a .roi file
//that can be loaded into the LAS AF microscope software. To do so, the macro will perform a tiling process when the design image (the binary image loaded)
//is bigger than the microscope tile (area that can be visualized at once with the micrscope). For each tile, the macro will detect all the particles in it. 
//The macro will then sort and group thee rois before writting their information to generate the .roi file.
//The files will be named as follow: roiName_z-coord_y-coord_x-coord. The coordinates correspond to the positions of the tile in the full image design.
//The input has to be a binary image with the LUT non-inverted, the area with value 255 (white) will be converted into rois.
//The macro will check if the image fulfill the requirements, if it does not, the macro will stop and the user has to modify the image.
//The grouping phase is needed to get the fewest groups possible so that they can be handled in the microscope.
//To perform the grouping process, the macro checks if a roi is included in ahother roi from the same group, 
//just in its boundning rectangle, or if they are independent. 
//The macro will ask the user to introduce the name and destination of the generated file, the size in microns of the microscope tile,
//the overlap between adjacent tiles and the design scale.
//Currently the macro just tolerate squared pixels and a maximun number of 10000 rois per tile. If more rois are wanted to be processed,
//it is necessary to lower the tile size or to modify the code at line 197 so that it includes more digits.
//The outputs of the macro will be stored in a foulder in the specified directory. It will include a .txt file containing the selected settings, 
//a reduced image of the design and the .roi files that can be loaded into the microscope. 
//A user guide for using this macro is available
if (is("Inverting LUT"))
{
	exit("The design has an inverting LUT. Please, invert it.")
}
designWidthPixels=getWidth();//Size in pixels of the full design loaded in ImageJ in pixels.
designHeightPixels=getHeight();
getStatistics(area, mean, min, max, std, histogram);
if (histogram[0]+histogram[255]!= (designWidthPixels*designHeightPixels))
{
	exit("The design is not binary. Please, binarize it.")
}
run("Set Measurements...", "area bounding redirect=None decimal=3");//Set the wanted measurements.
defName=getInfo("image.filename");//name of the design.
defDirectory=getInfo("image.directory");//directory of the design.
//GET THE INFORMATION OF THE DESTINATION SITE.
correctInfo=false;
while (correctInfo==false)//loop that controls that the introduced information is valid.
{
	Dialog.create("RoiLoader");//Dialog's box creation.
	Dialog.addMessage("Set file information:");
	Dialog.addString("Saving directory:\n",defDirectory,40);//Directory to save the results.
	Dialog.addCheckbox("Select a different directory (A new window will be opened after clicking ok)", false);//If selected (true), will display a get directory window.
	Dialog.addMessage("");
	Dialog.addString("Name:", defName, 40);//Ask for the name of the file that is going to be generated.
	Dialog.addMessage("Make sure that the name does not include spetial characters (/, \\, >, <, ?, *, :, |, \")\n----------------------------------------------------------------------------------------------------------\n");
	Dialog.addMessage("\nSet the design scale:");
	Dialog.addNumber("Number of pixels: ",5,0,9,"pixels");//Used to set the scale of the design.
	Dialog.addNumber("Expected size: ",5,0,9,"microns");
	Dialog.addMessage("----------------------------------------------------------------------------------------------------------\n");
	Dialog.addMessage("\nSet the tile size:");
	Dialog.addNumber("Tile width: ", 1550, 2, 9, "microns");//Ask for the size of each tile.
	Dialog.addNumber("Tile height: ", 1550, 2, 9, "microns");
	Dialog.addNumber("Overlap width: ", 0, 2, 9, "microns");//Ask for the overlap size of each tile.
	Dialog.addNumber("Overlap height: ", 0, 2, 9, "microns");
	Dialog.addMessage("----------------------------------------------------------------------------------------------------------\n");
	Dialog.addMessage("\nWrite any extra comments (Optional)");//Ask for any extra comments that will be writen in the .txt file.
	Dialog.addString("","",100);
	Dialog.show;
	//Dialog's box values retrieving.
savingDir=Dialog.getString();
changeSavingDir=Dialog.getCheckbox();
resultsName=Dialog.getString();
resultsName=split(resultsName,".");//Remove the part of the name after a dot (.), it will avoid mistakes when using names with extension included.
resultsName=resultsName[0];
numberOfPixels=Dialog.getNumber();
expectedSizeMicrons=Dialog.getNumber();
tileSizeWidthMicrons=Dialog.getNumber();
tileSizHeightMicrons=Dialog.getNumber();
overlapXMicrons=Dialog.getNumber();
overlapYMicrons=Dialog.getNumber();
extraComments=Dialog.getString();
//Error checking.
if ((numberOfPixels<=0)||(expectedSizeMicrons<=0)||isNaN(numberOfPixels)||isNaN(expectedSizeMicrons))//need to be a positive integrer.
{
	showMessage("Invalid scale. Please, reenter the data.");
}
else if ((tileSizeWidthMicrons<=0)||(tileSizHeightMicrons<=0)||isNaN(tileSizeWidthMicrons)||isNaN(tileSizHeightMicrons))//need to be a positive integrer.
{
	showMessage("Invalid tile size. Please, reenter the data.");
}
else if ((overlapXMicrons<0)||(overlapYMicrons<0)||isNaN(overlapXMicrons)||isNaN(overlapYMicrons))//need to be a positive or zero integrer.
{
	showMessage("Invalid overlap. Please, reenter the data.");
}
else
{
	correctInfo=true;//if everything is correct, goes out of the loop.
}
}//while correctInfo==false
//start = getTime(); 
//If another directory is wanted, display a get directory screen. Otherway, ensure that the directory name ends with /
if (changeSavingDir==true)
{
	savingDir = getDirectory("Choose a Directory in which to save the results");//Select a directory to save the generated file
}
else if (!endsWith(savingDir, "\\"))
{
	savingDir=savingDir+"\\";
}
//Create a folder to save all the results.
savingDirFinal=savingDir+resultsName+File.separator;//The name of the folder will be the same as the name of the file without the coordinates.
File.makeDirectory(savingDirFinal);
//Convert microns to meters.
expectedSizeMeters=expectedSizeMicrons*pow(10,-6);
tileSizeWidthMeters=tileSizeWidthMicrons*pow(10,-6);
tileSizeHeightMeters=tileSizHeightMicrons*pow(10,-6);
overlapXMeters=overlapXMicrons*pow(10,-6);
overlapYMeters=overlapYMicrons*pow(10,-6);
pixelSizeMicrons=expectedSizeMicrons/numberOfPixels;//Get pixel size.
pixelSize=pixelSizeMicrons*pow(10,-6);
tileWidthPixels=tileSizeWidthMeters/pixelSize;//Obtain the pixels of each tile according to the pixel size.
tileHeightPixels=tileSizeHeightMeters/pixelSize;
overlapX=overlapXMeters/pixelSize; //Overlap in pixels
overlapY=overlapYMeters/pixelSize;
//DUPLICATE AND CHECK IF THE design IS BINARY
run("Duplicate...", "title=temp_image duplicate");//Duplicate the design
selectWindow("temp_image");
run("8-bit");//Ensure that the design is 8-bit.
run("Invert","stack");
print("\\Clear");//clear log window
roiManager("reset");//Reset the RoiManager 
setBatchMode(true);
//TILING PROCESS:
//Calculate how many tiles must be done to cover the full design.
//Note that the designs must be rotated, so design width must match with tile height.
xSteps = designWidthPixels/(tileHeightPixels-overlapY);//number of times the tile photo will be included in the loaded design.
ySteps = designHeightPixels/(tileWidthPixels-overlapX);
if (floor(xSteps)==xSteps)//correct the value if there is not a natural number of tiles.
{
	roundUpXSteps=xSteps;
}
else
{
	roundUpXSteps=floor(xSteps)+1;
}
if (floor(ySteps)==ySteps)
{
	roundUpYSteps=ySteps;
}
else
{
	roundUpYSteps=floor(ySteps)+1;
}
/*
 * The next code will make a rectangle with a size corresponding to the tile size (tileHeightPixels and tileWidthPixels)
Each rectangle will generate an individual .roi file containing the rois included in the selected area. 
Finally the rectangle will move to the next tile to get the full design analyzed.
*/
setForegroundColor(255, 255, 255);//Select white as foreground color in order to properly draw the rectangles
totalXSizePixels=(tileHeightPixels-overlapY)*roundUpXSteps;
totalYSizePixels=(tileWidthPixels-overlapX)*roundUpYSteps;
run("Canvas Size...", "width=totalXSizePixels height=totalYSizePixels position=Center ");
for (stack=0; stack<nSlices;stack++)//for every stack
{
	setSlice(stack+1);
	if (nSlices>1)//If more than one slice, create a folder for each slice
	{
		savingDirFinalWithSlice=savingDirFinal+"z"+stack+File.separator;
		File.makeDirectory(savingDirFinalWithSlice);
	}
	else
	{
		savingDirFinalWithSlice=savingDirFinal;
	}
	for(k=0; k<roundUpXSteps; k++)//tile position in x
	{
		for(l=0; l<roundUpYSteps; l++)//tile position in y
		{
			roiManager("reset");//reset the roiManager
			posx=k+1;//Position of the tile with respect to the full design.
			posy=l+1;
			xIni=k*(tileHeightPixels-overlapY);//starting coordinates of the rectangle taking into account the overlap
			yIni=l*(tileWidthPixels-overlapX);
			makeRectangle(xIni, yIni, tileHeightPixels, tileWidthPixels); //make the rectangle (x_Initial, y_Initial, x_size, y_size)
			run("Duplicate...", "title=tile");//Duplicate the design of the corresponding tile
			selectWindow("tile");
			run("Rotate 90 Degrees Left");//rotate the design counterclockwise to overcome the rotation produced by the microscope
			tileWidth=getWidth();//size of the tile in pixels
			tileHeight=getHeight();
			makeRectangle(0, 0, tileWidth-1, tileHeight-1);
			run("Draw");//Draw the rectangle corresponding to the tile, that allow a correct Analyze Particles function when some item in the design is included in more than one tile.
			run("Select None");
			run("Outline");//Get the outline of the figures so that the Analyze Particles can detect inner particles.
			run("Analyze Particles...", "clear add");//Distinguish every Roi in the design and store them into the RoiManager.
			print("\\Clear");//clear the log window.
			{			
			//ORDER THE ROIS by size of their bounding rectangle, from bigger to smaller.
			run("Set Measurements...", "area bounding redirect=None decimal=3");
			boundingRectArea=newArray(roiManager("count"));//array that will contain the rank of the roi according to the area of the bounding rectangle
			for(i=0; i<roiManager("count");i++)
			{
				roiManager("select",i);
				run("Measure");
				area=getResult("Width",i)*getResult("Height",i);
				boundingRectArea[i]=area; //store the areas
			}
			rankedBoundingRectArea=Array.rankPositions(boundingRectArea);//sort the areas.
			reversedRankedBoundingRectArea=Array.reverse(rankedBoundingRectArea);//the rois are ordered from smaller to bigger, so we invert it.
			for(i=0; i<roiManager("count");i++)//rename the rois according to their bounding rectangle area rank.
			{
				roiManager("select", reversedRankedBoundingRectArea[i]);
				if (i<10)
				{
					roiManager("rename", "000"+i); //the 0s are added in order to make the sort, as the names are considered as strings.
				}
				else if (i<100)
				{
					roiManager("rename", "00"+i);
				}
				else if (i<1000)
				{
					roiManager("rename", "0"+i);
				}
				else if (i<10000)
				{
					roiManager("rename", i);
				}
				else
				{
					selectWindow("tile");
					close();
					selectWindow("temp_image");
					close();
					selectWindow("ROI Manager");
					run("Close");
					selectWindow("Log");
					run("Close");
					selectWindow("Results");
					run("Close");
					exit("Too many rois detected.\nConsider a smaller tile size")//If there are too many rois, the macro will be aborted.
					//Add more else if to consider a larger number of rois  if needed.
				}
			}
			roiManager("sort");
			/*The rois are ordered by the size of their bounding rectangle			
			 * GROUP THE ROIS: starting with the biggest, change the name to group-index.
			It will place the rois in the same group if they are not included in any of the bounding rectangles of the other rois in the group
			or if they are included just in one of the other rois in the group.
			*/
			changes=true;//Boolean variable that will get true if any name is changed in the Rois Manager.
			group=-1;//Number of groups found. It starts in -1 because the last turn of the loop will not make any change.
			roisGrouped=newArray(roiManager("count"));//Array that will contain the indexes of the rois included in the current group. Also will restart the variable.
			while (changes==true) //will get out of the loop if no changes are made in the names of the rois, when all the rois are properly named.
			{
				group++;//Number of groups found.
				changes=false;//Will change into true if any change is made.
				roisInGroup=0;//Number of Rois in the current group.
				for (i=0;i<roiManager("count");i++)//For all the roi manager.
				{
					roiManager("select",i);
					in=0;//Will be used to group the rois.
					if ((! matches(selectionName,".*-.*")))//the grouped rois will have the format group-index, so if the name does not have a - it is not grouped yet
					{
						if (roisInGroup==0)//for the first roi in each group:
						{
							roiManager("rename",group+"-"+i); //Rename
							changes=true;
							roisGrouped[roisInGroup]=i; //Store the index of the grouped roi in an array. 
							roisInGroup++;
						}
						else //If it is not the first roi in the group.
						{
							/*For all the rois that are already in the group (j), check if the next one (i) in included in any of them							
							 * 
							or in their bounding rectangle.
							*/
							for (j=0;j<roisInGroup;j++)
							{
								roiManager("Select",i);
								getSelectionBounds(xIni, yIni, widthRect, heightRect);//Will return the upper left corner of the bounding rectangle containing the selection and its width and height
								//Get the vertex of the bounding rectangle
								coord1=newArray(2);
								coord1[0]=xIni;
								coord1[1]=yIni;
								coord2=newArray(2);
								coord2[0]=xIni+widthRect-1;//correct the outline shift
								coord2[1]=yIni;
								coord3=newArray(2);
								coord3[0]=xIni;
								coord3[1]=yIni+heightRect-1;//correct the outline shift
								coord4=newArray(2);
								coord4[0]=xIni+widthRect-1;//correct the outline shift
								coord4[1]=yIni+heightRect-1;//correct the outline shift
								Roi.getCoordinates(xpoint, ypoint);//Get the coordinates of the roi
								roiManager("Select",roisGrouped[j]);
								//print(coord1[0],coord1[1],coord2[0],coord2[1]);
								if ((selectionContains(xpoint[0],ypoint[0])==1))//if it is included in the roi
								{
									in++;//Number of rois it is cointained in.
									
									//print (xpoint[0],ypoint[0]);
									//waitForUser("in++");
								}
								else if (in==0) //If it is not included in any rois.
								{
									getSelectionBounds(xIni, yIni, widthRect, heightRect);//Select the bounding rectangle.
									makeRectangle(xIni, yIni, widthRect, heightRect); 
									if ((selectionContains(coord1[0],coord1[1])==1)&&(selectionContains(coord2[0],coord2[1])==1)&&
									(selectionContains(coord3[0],coord3[1])==1)&&(selectionContains(coord4[0],coord4[1])==1))//If it is included in the bounding rectangle
									{
										in=2;
										//waitForUser("in2");
									}
								}//else
							}//For roisIngroup
							//print (in);
							//waitForUser("");
							if (in<=1)//If it is not included in any bounding rectangle (in=0) or if it is included in just one roi (in=1) change the name to include it in the group.
							{
								roiManager("select",i);
								roiManager("rename",group+"-"+i);
								changes=true;
								roisGrouped[roisInGroup]=i; //Add it to the array, so the next roi will be checked against it.
								roisInGroup++;
							}//if in<=1	
						}//else: roinsingroup=~0
					}//if Matches
				}//for roimanager Count
			}//while changes
			//waitForUser("");
			/*ALL ROIS ARE GROUPED IN THE ROI MANAGER			
			Now the macro will print the .roi format, making a roi Composite (group of rois grouped that can be handle as one in LasAF) for every previously generated group.
			*/
			//Print the opening of the .roi file that is going to be generated.
			print("<LMSDataContainerHeader Version=\"2\">");
			print("<Element Name=\"Save\">");
			print("<Data>");
			print("<ROISet ROISetType=\"2\" PossibleChildROITypes=\"25394\" PossibleROITransforms=\"65535\" PossibleROIActions=\"19\" />");
			print("</Data>");
			print("<Memory Size=\"0\" MemoryBlockID=\"MemBlock_268\"/>");
			print("<Children>");
			for (groupCounter=0;groupCounter<group;groupCounter++)
			{
				roisInGroup2=newArray(roiManager("count"));//Array that will contain the indexes of the rois included in the group.
				roisNumber=-1;//Number of rois in each group.                 	
				for (roiCounter=0;roiCounter<roiManager("count");roiCounter++)//For each roi. 
				{
					//If roi in pos roiCounter is in the group groupCounter, store the indexes in an array
					roiManager("select",roiCounter);
					if(matches(selectionName,".*"+groupCounter+"-.*"))//The name is ***group-***
					{
						roisNumber++;
						roisInGroup2[roisNumber]=roiCounter;
					}
				}//for roiCounter
				rois=Array.trim(roisInGroup2, roisNumber+1);//take only the useful information in roisInGroup2, that will be the indexes of the rois in that group
				//Combine all the rois in the group in order to obtain its center. This is needed to get the correct position of the group
				roiManager("Select",rois);
				if(lengthOf(rois)>1)
				{
					roiManager("Combine");
				}
				getSelectionBounds(xIni, yIni, widthRect, heightRect);//Bounding rectangle of the combined rois
				groupPosX=(xIni+xIni+widthRect)/2;//Center of the group
				groupPosY=(yIni+yIni+heightRect)/2;
				convertedGroupPosX=(groupPosX*pixelSize)-(0.5*tileSizeWidthMeters);//Conversion of the center of the group from pixels to meters and translation to LAS AF origin of coordinates
				convertedGroupPosY=(groupPosY*pixelSize)-(0.5*tileSizeHeightMeters);
				//PRINT THE INFORMATION FOR EACH GROUP
				print("<Element Name=\"\">");
				print("<Data>");
				print("<ROIComposite RoiType=\"56\" RoiAction=\"65535\" TransformationType=\"65535\" Color=\"4294901760\" >");
				print("<Transformation Rotation=\"0\">");
				print("<Scaling XScale=\"1\" YScale=\"1\"/>");
				print("<Translation X=\"",convertedGroupPosX,"\" Y=\"",convertedGroupPosY,"\"/>");//write the translation of the group.
				print("</Transformation>");
				print("</ROIComposite>");
				print("</Data>");
				print("<Memory Size=\"0\" MemoryBlockID=\"MemBlock_268\" />");
				print("<Children>");
				/*Group Print is completed.                        	
				PRINT THE INFORMATION FOR EACH ROI IN THE GROUP
				*/
				for (roiCounter=0;roiCounter<lengthOf(rois);roiCounter++)
				{
					roiManager("deselect");
					roiManager("select",rois[roiCounter]);//Select the roi in the group
					print("<Element Name=\"\">");
					print("<Data>");
					print("<ROISingle RoiType=\"8\" RoiAction=\"65535\" TransformationType=\"65535\" Color=\"4294901760\" IsClosed=\"1\" Inverted=\"0\">");
					print("<Vertices>");
					Roi.getCoordinates(xpoints, ypoints);//Obtain the coordinates of the roi.
					for(j=0;j<lengthOf(xpoints);j++)//Transform and print the coordinates.
					{
						conversionOfXCoord=((xpoints[j])*pixelSize)-(0.5*tileSizeWidthMeters);//Transformation of the coordinates.
						conversionOfYCoord=((ypoints[j])*pixelSize)-(0.5*tileSizeHeightMeters);
						//Print the coordinates with the appropriate  format.
						print("<P X=\"",conversionOfXCoord,"\" Y= \"",conversionOfYCoord,"\"/>");
					}
					//Print the .roi format for the end of each roi.
					print("</Vertices>");
					print("<Transformation Rotation=\"0\">");
					print("<Scaling XScale=\"1\" YScale=\"1\"/>");
					print("<Translation X=\"0\" Y=\"0\"/>");
					print("</Transformation>");
					print("</ROISingle>");
					print("</Data>");
					print("<Memory Size=\"0\" MemoryBlockID=\"MemBlock_268\" />");
					print("<Children />");
					print("</Element>");
				}//for roiCounter.
				//print the .roi format for the end of each group.
				print("</Children>");
				print("</Element>");
			}//group.
			//print the .roi format for the end of the file.
			print("</Children>");
			print("</Element>");
			print("</LMSDataContainerHeader>");
			//save the file and change the extension from .txt to .roi.
			selectWindow("Log");
			if (roiManager("count")==0)	
			{
				selectWindow("Log");
				saveAs("text",savingDirFinalWithSlice+resultsName+"_z"+stack+"_y"+posy+"_x"+posx+"Empty"+".txt");
				path1= savingDirFinalWithSlice+resultsName+"_z"+stack+"_y"+posy+"_x"+posx+"Empty"+".txt";
				path2= savingDirFinalWithSlice+resultsName+"_z"+stack+"_y"+posy+"_x"+posx+"Empty"+".roi";
			}
			else if	((roiManager("count")==1)&&(getResult("Area",0)==((tileWidth-2)*(tileHeight-2))))//if there is just one roi that correspond to the tile, it is considered an empty tile.
			{
				saveAs("text",savingDirFinalWithSlice+resultsName+"_z"+stack+"_y"+posy+"_x"+posx+"Full"+".txt");
				path1= savingDirFinalWithSlice+resultsName+"_z"+stack+"_y"+posy+"_x"+posx+"Full"+".txt";
				path2= savingDirFinalWithSlice+resultsName+"_z"+stack+"_y"+posy+"_x"+posx+"Full"+".roi";
			}
			else
			{
				saveAs("text",savingDirFinalWithSlice+resultsName+"_z"+stack+"_y"+posy+"_x"+posx+".txt");
				path1= savingDirFinalWithSlice+resultsName+"_z"+stack+"_y"+posy+"_x"+posx+".txt";
				path2=savingDirFinalWithSlice+resultsName+"_z"+stack+"_y"+posy+"_x"+posx+".roi";
			}
			File.rename(path1, path2);
			//waitForUser("");
			selectWindow("tile");
			close();
		}//if roimanager==0
		}//for l.
	}//for k.
}//for stack
selectWindow("temp_image");
close();
selectWindow("ROI Manager");
run("Close");
selectWindow("Log");
run("Close");
selectWindow("Results");
run("Close");
//Write the .txt file.
print("File name:",resultsName,"\nScale:\nNumber of pixels:",numberOfPixels,"\nExpected size:",expectedSizeMicrons,"\n\tPixel size (um/pixel):"
,pixelSizeMicrons,"\nTile width (um):",tileSizeWidthMicrons,"\nTile height (um):",tileSizHeightMicrons,"\nOverlap width (um):"
,overlapXMicrons,"\nOverlap height (um):",overlapYMicrons,"\nNumber of slices:",nSlices,"\nExtra comments:",extraComments); //Writting all the data corresponding to the setting and the extra comments
selectWindow("Log");
saveAs("text",savingDirFinal+resultsName+"_Log.txt"); //save the log window as a .txt file.
run("Close");
//resize the design and perform transformations so that it can be stroed in the foulder with minimun size.
selectWindow(defName);
run("Duplicate...", "title="+defName+" duplicate");
//reduce the longest dimension to 512 pixels
if ((designHeightPixels>=designWidthPixels)&&(designHeightPixels>512))
{
	resizedHeight=512;
	resizedWidth=designWidthPixels*resizedHeight/designHeightPixels;
}
else if ((designHeightPixels<=designWidthPixels)&&(designWidthPixels>512))
{
	resizedWidth=512;
	resizedHeight=designHeightPixels*resizedWidth/designWidthPixels;
}
else
{
	resizedHeight=designHeightPixels;
	resizedWidth=designWidthPixels;
}
run("Size...", "width="+resizedWidth+" height="+resizedHeight+" depth="+nSlices+" constrain average interpolation=Bilinear"); //re-scale
run("8-bit");//transform the design into 8-bit
//run("Make Binary");//make binary
//setOption("BlackBackground", false);
saveAs("tiff",savingDirFinal+resultsName+".tif"); //save the design
//print((getTime()-start)/1000);  
showMessage("End of process, your results are available at:\n"+savingDirFinal);
```
